# Analysis of isobaric quantitative proteomic data using TMT-Integrator and FragPipe computational platform

**DOI:** 10.1101/2025.05.27.656447

**Authors:** Hui-Yin Chang, Yamei Deng, Ruohong Li, Dmitry Avtonomov, Bo Wen, Sarah E. Haynes, Felipe da Veiga Leprevost, Bing Zhang, Fengchao Yu, Alexey I. Nesvizhskii

**Affiliations:** Department of Pathology, University of Michigan, Ann Arbor, MI, USA; Department of Biomedical Sciences and Engineering, Institute of Systems Biology and Bioinformatics, National Central University, Taiwan; Lester and Sue Smith Breast Center, Department of Molecular and Human Genetics, Baylor College of Medicine, One Baylor Plaza, Houston, TX 77030, USA; Gilbert S. Omenn Department of Computational Medicine and Bioinformatics, University of Michigan, Ann Arbor, MI, USA

## Abstract

Isobaric mass tags, such as iTRAQ and TMT, are widely utilized for peptide and protein quantification in multiplex quantitative proteomics. We present TMT-Integrator, a bioinformatics tool for processing quantitation results from TMT and iTRAQ experiments, offering integrative reports at the gene, protein, peptide, and post-translational modification site levels. We demonstrate the versatility of TMT-Integrator using five publicly available TMT datasets: an *E. coli* dataset with 13 spike-in proteins, the clear cell renal cell carcinoma (ccRCC) whole proteome and phosphopeptide-enriched datasets from the Clinical Proteomic Tumor Analysis Consortium, and two human cell lysate datasets showcasing the latest advances with the Astral instrument and TMT 35-plex reagents. Integrated into the widely used FragPipe computational platform (https://fragpipe.nesvilab.org/), TMT-Integrator is a core component of TMT and iTRAQ data analysis workflows. We evaluate the FragPipe/TMT-Integrator analysis pipeline’s performance against MaxQuant and Proteome Discoverer with multiple benchmarks, facilitated by the bioinformatics tool OmicsEV. Our results show that FragPipe/TMT-Integrator quantifies more proteins in the E. coli and ccRCC whole proteome datasets, identifies more phosphorylated sites in the ccRCC phosphoproteome dataset, and delivers overall more robust quantification performance compared to other tools.

## INTRODUCTION

Tandem mass spectrometry (MS/MS) in combination with liquid chromatography (LC), LC-MS/MS, has become a central technology in the field of proteomics [1, 2]. Isobaric mass tags such as iTRAQ and TMT are isotopically-encoded chemical reagents composed of a mass reporter region (which determines the mass channel), a balancer region, and a reactive group that attaches to peptide N-termini and lysine residues [3–7]. In isobaric quantitative proteomics workflows, digested peptides from different biological samples are labeled with reagents from a different channel and then combined for LC-MS/MS analysis. During mass spectrometry analysis, the reagents are expected to dissociate such that the mass reporter regions from each channel can be distinguished in the MS/MS spectra. Peptide and protein quantification between samples/channels is then accomplished by comparing the reporter ion intensities within MS/MS spectra. Since the interference effect (e.g., co-fragmentation) inevitably exists during the acquisition of tandem mass spectra [8, 9], triple-stage mass spectrometry (MS3) has been proposed to mitigate potential ratio distortion [10–12]. Isobaric labeling enables simultaneous measurement of multiple samples, reducing analysis time and eliminating potential run-to-run variation [13]. The commercially available isobaric chemical tags facilitate the simultaneous analysis (multiplexing) of up to 18 (recently extended to 32) experimental samples [14]. The isobaric mass tag-based strategies are now routinely applied as part of biological studies, from drug target identification [15] to interactome profiling [16, 17] to single cell proteomics [18–20]. It has been extensively applied in multiple proteogenomic studies such as those supported by the Clinical Proteomic Tumor Analysis Consortium (CPTAC) [21–27].

There have been multiple publications describing various computational approaches developed specifically for post-processing and differential protein analysis of TMT/iTRAQ proteomics data [28–34], including MSstatsTMT [35] and PAW pipeline [36]These tools focus on improving the integration of quantitative data, acting as independent components to support isobaric quantification. Still, most isobaric mass tag-based proteomics studies so far used the Proteome Discoverer, largely because of its end-to-end analysis capabilities and a graphical user interface (GUI). However, at least in our own experience of working with CPTAC data, application of Proteome Discoverer, and other tools such as MaxQuant [37], to large datasets consisting of many TMT-labeled sample sets (plexes) has been challenging. Thus, we sought to create a new option for the proteomics community that would enable fast and accurate analysis of such data that can process datasets with dozens or even hundreds of TMT plexes, and that can be run on a variety of computational platforms, from individual Windows desktop to high performance Linux clusters to cloud computing. This resulted in the creation of a new computational platform, FragPipe, that included our ultrafast database search engine MSFragger [38, 39], the Philosopher toolkit [40], quantification module IonQuant [41, 42], and multiple other tools for comprehensive analysis of proteomics data, executable via command line or an easy-to-use GUI. TMT-Integrator has been a key component of the pipeline, enabling aggregation of PSM tables containing raw TMT reported ion intensities extracted by IonQuant or Philosopher from one or more TMT plexes experiment, followed by PSM to peptide and protein-level roll-up and report generation. TMT-Integrator not only leverages abundance information from peptide-spectrum matches (PSMs) to higher-level quantification (e.g., modified site, peptide, protein, and gene) but also supports several outlier removal and normalization options for downstream statistical analyses. We have used TMT-Integrator for all proteomics and phosphoproteomics analyses as part of multiple CPTAC landmark publications [21, 22, 24, 25, 43], and other studies [44].

Here, we present a more in-depth description of TMT-Integrator, as well as evaluate the performance of our entire analysis pipeline using five proteomics datasets. The first two are the whole proteome and phosphopeptide-enriched datasets from the clear cell renal cell carcinoma (ccRCC) study published by the CPTAC [21]. Each of the two datasets consists of 23 TMT 10-plexes of patient tumor and adjacent normal tissue samples, with replicates of two additional samples (a non-CPTAC kidney tumor sample and an NCI-7 Cell Line mix sample) inserted in the dataset for quality control. The third dataset is a single 10-plex *E. coli* experiment [45] downloaded from the ProteomeXchange Consortium [46] website (with identifier PXD005486), where 13 standard proteins were spiked in at different amounts. The other two datasets contain human cell lysates derived from diverse cell lines in biological triplicates. One is collected on an Astral mass spectrometer, while the other utilizes the newly developed TMT 35-plex reagents which enable simultaneously analyzing up to 35 samples. The five datasets were processed using built-in FragPipe TMT workflows, and the identification and quantification results were evaluated using OmicsEV [47]. According to the analyses of ccRCC whole proteome data, our pipeline identifies more proteins than MaxQuant and exhibits reduced batch effects and higher gene-wise correlation matched with mRNA data. FragPipe with TMT-Integrator has similar performance to Proteome Discoverer and MaxQuant computational platforms in detecting spike-in proteins in the *E. coli* sample. We also demonstrate that FragPipe with TMT-Integrator performs well for analyzing Astral and TMT 35-plex data while ensuring precise and accurate quantification. In addition to providing accurate quantification, TMT-Integrator can be easily applied to a range of experimental designs, from individual single-shot samples to large-scale studies with many multiplexed samples and fractions.

## RESULTS

### Overview of the isobaric quantification workflow in FragPipe

The overview of the entire TMT data analysis workflow in FragPipe is shown in **Figure 1**. It consists of three main steps: (1) peptide identification from MS/MS spectra using MSFragger; (2) PSM validation, protein inference, FDR filtering, and quantification using various FragPipe tools; (3) PSM table integration, analysis, and report generation using TMT-Integrator. The output files from MSFragger (pepXML files) are processed using Percolator [52] (alternatively PeptideProphet [64]) to compute the posterior probability of correct identification for each PSM. The output from PeptideProphet (pep.xml files) or from Percolator (after conversion of its output to pep.xml format) from all files in all TMT plexes in the experiment are then processed with ProteinProphet [54] together to assemble peptides into proteins (protein inference) and to create a combined file (in prot.xml format) of high confidence protein groups. In the case of PTM-enriched datasets, Percolator (or PeptideProphet) output pep.xml files are additionally processed using PTMProphet [53] to calculate the localization probabilities for each modification site (generating modified, in mod.pep.xml format, files). This combined protXML file from ProteinProphet, and the individual pep.xml (or mod.pep.xml) files for each TMT plex are further processed using the Philosopher filter module. As part of this step, each peptide is assigned either as a unique peptide to a particular protein or assigned as a razor peptide to a single protein with the most peptide evidence. When two or more proteins are indistinguishable given the set of observed peptides, one entry is selected as representative protein for that indistinguishable protein group. The protein list is filtered to 1% protein-level False Discovery Rate (FDR) using the best peptide approach (considering both unique and razor peptides). By default, the picked FDR target-decoy strategy is used in all TMT workflows [65]. In each TMT plex, the PSM lists are filtered using a sequential FDR strategy, retaining only those PSMs that pass TMT set-specific 1% PSM-level FDR and map to proteins that also passed the global 1% protein-level FDR filter. For each PSM that passes these filters, the corresponding precursor ion MS1 intensity is extracted using IonQuant (alternatively, as used in earlier versions of the pipeline, using the label-free quantification module of Philosopher). Finally, for every PSM corresponding to a TMT-labeled peptide, TMT reporter ion intensities are extracted from the MS/MS scans using IonQuant (alternatively, using the Philosopher’s isobaric quantification command), which also calculates precursor ion purity scores. All supporting information for each PSM, including the accession numbers and names of the protein/gene selected based on the protein inference approach with razor peptide assignment and quantification information (MS1 precursor-ion intensity and the TMT reporter ion intensities), are summarized in the output PSM.tsv files, one file for each TMT plex. TMT-Integrator takes PSM tables (one table for each TMT plex) as input and exports an integrated report with columns for sample names and rows for abundances at each user-specified level of data summarization (gene, protein, peptide, PTM multi-site, and PTM single-site). The key steps of the TMT-Integrator algorithms (see **Methods**) include additional filtering of PSMs, reference normalization (conversion of PSM-level reporter ion intensities to ratios), grouping of PSMs at each level, outlier removal, PSM abundance aggregation (i.e., quantification roll-up from PSMs to each higher data summarization level), sample-wise (across all peptides/proteins) ratio normalization, and finally conversion from ratios back to absolute intensities (“abundances”). At the end, TMT-Integrator prints multiple output tables, containing ratios (to the reference sample, or to internally created virtual reference; ‘ratio’ labeled tables) and absolute intensities (‘abundance’ labeled tables). The tables are additionally labeled to indicate the level of data summarization (gene, protein, peptide, multi-site, single-site) and the normalization method (None, MD, GN), e.g., abundance_gene_MD.tsv, ratio_protein_None.tsv, etc.

**Figure 1.**
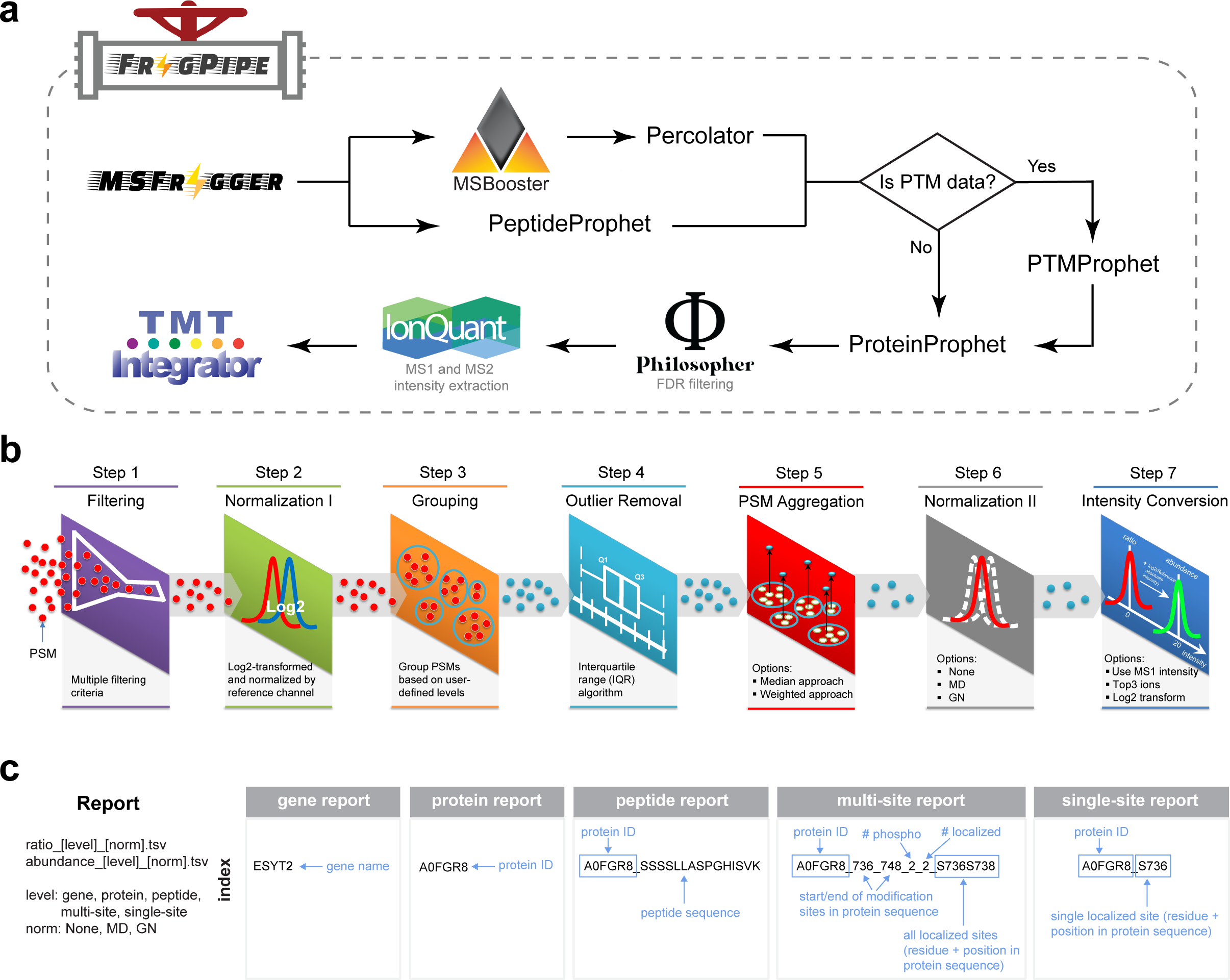
Overview of isobaric labeling data analysis with TMT-Integrator in FragPipe. **(a)** FragPipe workflow for isobaric labeling data analysis in both global proteomic data and post-translational modification (PTM)-enriched proteomic data, with PTMProphet used specifically in PTM data analysis. **(b)** Seven major steps included in TMT-Integrator. **(c)** Summary of TMT-Integrator report files and the breakdown of index formats across report levels, illustrated with an example.

### Ratio-based integration for analyzing samples from multiple plex sets

Due to the limited multiplexity of isobaric labeling reagents for simultaneously labeling and analyzing samples in a single MS experiment, multiple plex sets are usually required to analyze samples in a scale that exceeds its multiplexity, and a channel in each plex set is commonly used as the reference channel, typically containing equal amount of protein from pooled samples. This common reference channel helps bridge multiple plex sets and is used to normalize the data to minimize the batch effect. However, a real reference sample may not always be included. To support both scenarios, we implemented two approaches in TMT-Integrator to perform ratio-to-reference normalization, as shown in **Figure 2a**. One is the real reference approach for data with real reference channels, and the other is a virtual approach that involves creating a virtual reference channel by taking a median of intensities across each plex. Ratio-to-reference normalization is then performed against either the real or virtual reference channel for each plex. The virtual approach is necessary when a separate reference sample has not been included (as is often the case in experiments involving a single TMT plex), and it can also facilitate combining data from different cohorts or studies.

**Figure 2.**
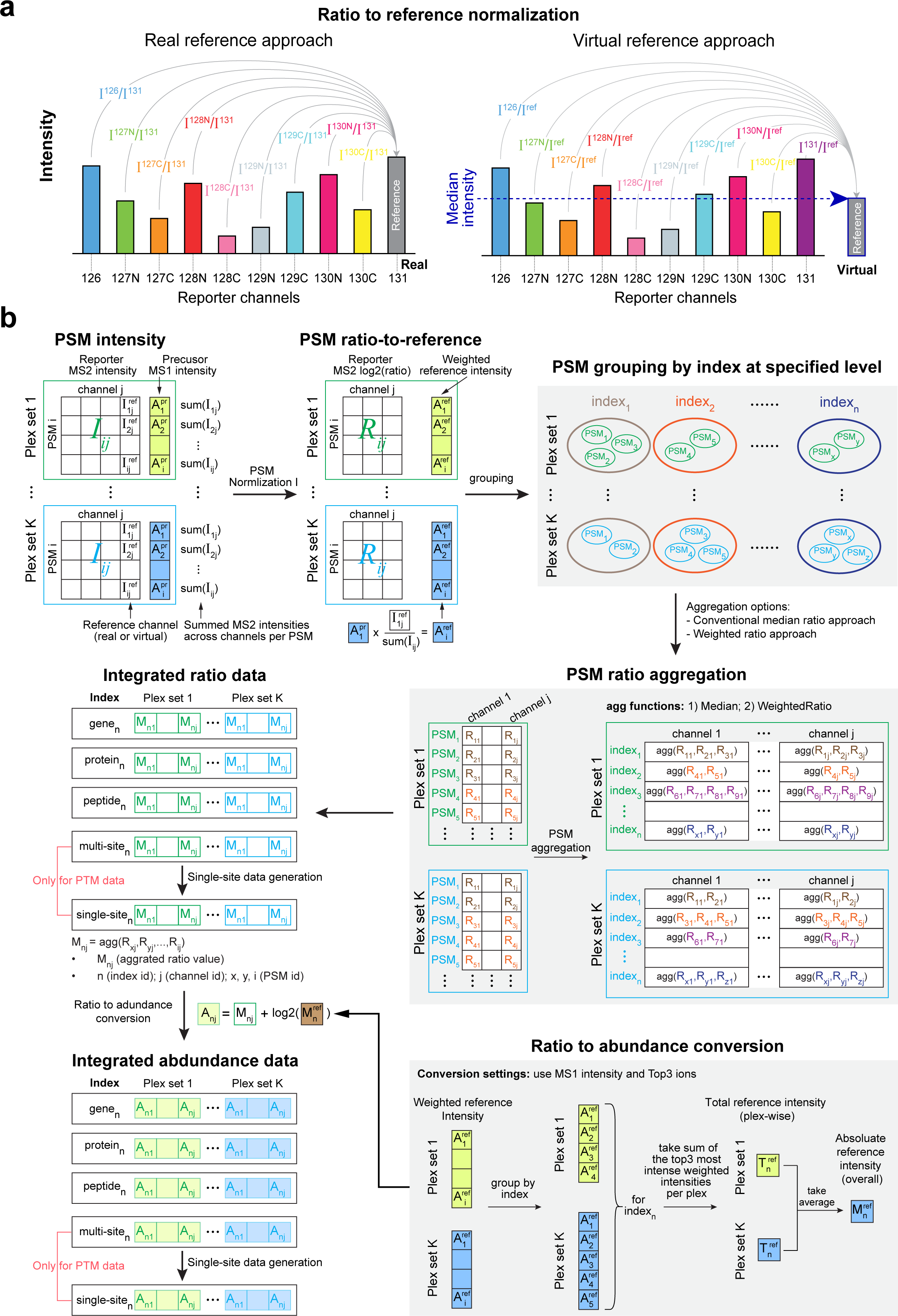
Illustrations of PSM ratio integration in TMT-Integrator across multiple plex sets. **(a)** Two approaches for PSM ratio-to-reference normalization in Normalization I. Using the TMT 10-plex for illustration, each bar represents a channel, and its height represents the measured intensity. **(b)** Schematic diagram illustrating how TMT-Integrator processes a multi-plex dataset in three main steps: (1) transforming the PSM intensity table to a PSM ratio-to-reference table through normalization, (2) integrating PSM ratio tables from multiple plex sets into a single ratio matrix at a specific level, and (3) converting the ratio matrix into an abundance matrix by incorporating absolute intensities.

With the help of the ratio-to-reference normalization, TMT-Integrator can efficiently integrate data from multi-plex experiments. To provide a comprehensive overview of this integration process, we therefore break down the whole process into four major steps primarily from the quantitative data transformation perspective, including : (1) transforming the PSM intensity table to a PSM ratio-to-reference table through normalization, (2) integrating PSM ratio tables from multiple plexes into a single ratio matrix at a specific level, and (3) converting the ratio matrix into an abundance matrix by incorporating absolute intensities. For PTM data, an additional step generates single-site information from multi-site data (see **Figure 2b**). TMT-Integrator starts by taking the PSM tables as input. TMT-Integrator extracts MS2 intensities for reporter channels and MS1 intensities for precursors from PSM.tsv files to create the PSM intensity table, where reporter intensity values are denoted as I_ij_ (i=PSM ID, j=Channel ID) and precursor intensity values are denoted as A^Pr^ . Channel intensities are normalized using the reference channel and log2-transformed to produce ratio tables for each plex, where intensity values are denoted as R_ij_ (i=PSM ID, j=Channel ID). At the same time, MS1 precursor abundances are transformed into reference-weighted abundances by multiplying them by a factor representing the proportion of the reference channel intensity in the total MS2 (sum across all channels) intensity, denoted as A^ref^_i_ (i=PSM ID). PSMs from each plex are further grouped by index at a specified level, and PSM ratios are then aggregated based on the grouping information. There are two options for conducting ratio integration: the conventional median ratio approach and the weighted ratio approach (see **Methods**). After aggregating PSM ratios to their respective indexes for each channel, the aggregated ratios from all plexes are combined into an integrated ratio table, where the ratio value is denoted as M_nj_ (n is the index ID and j is the channel ID). The ratio matrix is further converted to an abundance matrix by adding back absolute intensities of reference channel (denoted as M^ref^_n_) aggregated from reference-weighted abundances for each group (index_n_). In this matrix, abundance value is denoted as A_nj_ (n=index ID, j=channel ID). In contrast to using MS1 intensity and top3 most intense peptide ions to derive absolute intensities per group herein, using MS2 intensity and all peptide ions is also supported for ratio-to-abundance conversion. For PTM data, the single-site matrix is generated from the multi-site matrix (**Supplementary** Figure 1**; Methods**) by collapse multiple modified forms into singly modified sites by applying the following logic (applied subsequently): (1) keep only PSMs with localized sites; (2) use only PSMs with the single localized site when available; (3) use median of all localized sites if only found with other sites.

### Robust and accurate quantification by TMT-Integrator in large-scale datasets

We first evaluated the performance of FragPipe with TMT-Integrator alongside MaxQuant, using the ccRCC whole proteomic dataset, which has 23 TMT 10-plexes (each plex contains 25 fractions). To simplify the comparison between the tools and with matching RNA data, the analysis was performed using protein quantification data collapsed to the gene symbol level (see **Methods**). Both TMT-Integrator and MaxQuant were run with the normalization to actual reference sample (the pooled sample) as well as using the virtual reference approach, referred to as ‘FP/MQ RefRatio’ and ‘FP/MQ VirtualRatio’, respectively (**Table 1**). For both tools, unless otherwise noted, we used the conventional median approach when aggregating ratios-to-reference from PSM to peptide/protein level. An important difference between the two tools is that FragPipe performs an additional step of converting from ratios-to-reference back to absolute protein abundances, whereas MaxQuant produces the ratio tables only. For both tools, the output tables were computed with and without an additional median centering (‘MD’) of all protein quantifications in each sample. For MaxQuant, this median centering was applied using additional scripts (**Code Availability**, the default output from MaxQuant is not median centered). We also run MaxQuant using its “No Normalization” approach (‘MQ NoNorm’, and ‘MQ NoNorm MD’), in which no ratio to a reference sample (actual or virtual) is taken. Note that under this option – which is the default mode of MaxQuant - MaxQuant generates output tables with absolute peptide/protein intensities since no normalization to reference is performed.

**Table 1.** Evaluation of FragPipe with TMT-Integrator (FP) and MaxQuant (MQ) results using ccRCC whole proteomic dataset in terms of the number of proteins (genes), in total and with a detection rate above 50%. Also shown the Complex K-S statistic, and gene-wise and sample-wise abundance correlations between protein and RNA data (R_gene-wise_, R_sample-wise_). Additional global median centering normalization is indicated with MD (for MaxQuant, performed outside of MaxQuant).

| Method | Proteins<br>(< 50% missing) | $R_{\text{gene-wise}}$ | $R_{\text{sample-wise}}$ | complex_KS | func_auc |
| --- | --- | --- | --- | --- | --- |
| FP RefRatio | 12210 (9521) | 0.28 | 0.46 | 0.82 | 0.83 |
| FP RefRatio MD (default) | 12210 (9521) | 0.41 | 0.46 | 0.86 | 0.82 |
| FP VirtualRatio MD | 12210 (9521) | 0.37 | 0.46 | 0.87 | 0.83 |
| FP RefRatio Weighted MD | 12210 (9521) | 0.39 | 0.47 | 0.85 | 0.83 |
| MQ NoNorm (default) | 10815 (8463) | 0.19 | 0.41 | 0.65 | 0.65 |
| MQ NoNorm MD | 10815 (8463) | 0.21 | 0.41 | 0.62 | 0.70 |
| MQ RefRatio MD | 10815 (8463) | 0.41 | 0.19 | 0.84 | 0.72 |
| MQ VirtualRatio MD | 10815 (8463) | 0.37 | 0.17 | 0.86 | 0.74 |
| MQ VirtualRatio Weighted MD | 10815 (8463) | 0.24 | 0.29 | 0.71 | 0.73 |

The number of quantified proteins (in total, and with less than 50% missing values across the entire cohort) reported by TMT-Integrator was higher than that reported by MaxQuant (12210 vs. 10815 in total; 9521 vs. 8463 with less than 50% missing rate; **Table 1** and **Figure 3a**). Of note, 10556 proteins (85%) were commonly detected by both tools. Both tools were able to accurately separate tumor from normal samples (see **Figure 3b** for FragPipe and **Supplementary** Figure 2a for MaxQuant). Using default settings for each pipeline, FragPipe with TMT-Integrator (‘FP RefRatio MD’) and MaxQuant (‘MQ NoNorm’), both with absolute abundance intensities, showed similar protein abundance correlation between replicate runs of the same QC sample (5 replicate runs); however, FragPipe demonstrated better correlation across the replicates of the 7 NCI-7 sample runs (**Figure 3c**). The NCI-7 samples are different from the patient tissue samples and the QC control tissue sample. Thus, this observation suggests that TMT-Integrator with normalization to reference approach was better able to account for the outlier nature of the NCI7 samples in this cohort. We also compared protein abundance correlation using data tables with ratio-to-reference intensities: FragPipe ratio report with default settings (‘FP RefRatio MD’) and MaxQuant output with the normalization to reference settings alongside median-centering normalization (‘MQ RefRatio MD’). Both tools consistently showed overall good and similar correlations for NCI and QC samples, indicating their comparable performance in achieving reliable ratio-to-reference quantifications (see **Supplementary** Figure 2b).

**Figure 3.**
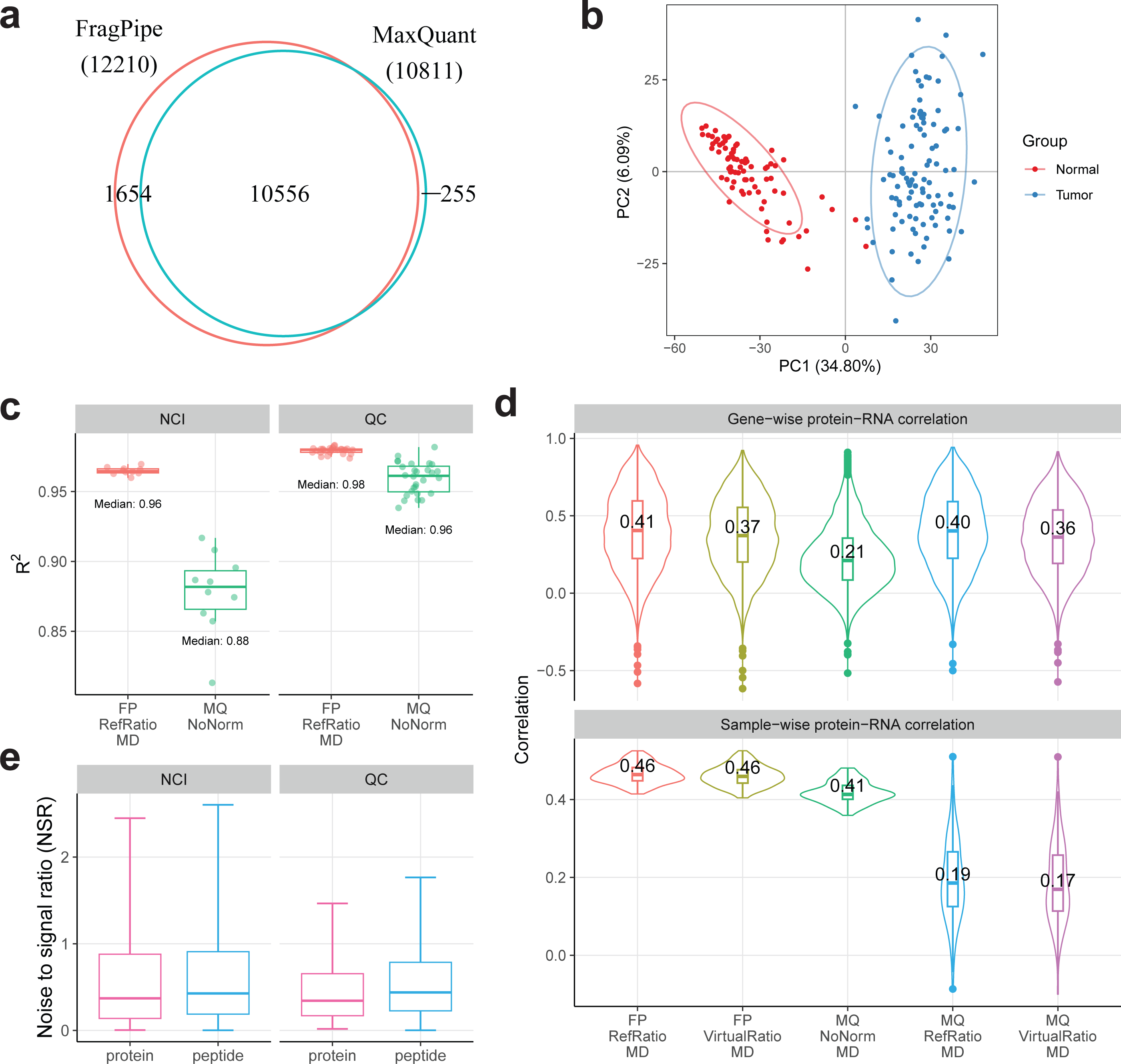
Performance evaluations on ccRCC whole proteome dataset. **(a)** Venn diagram showing the number of genes identified by FragPipe/TMT-Integrator and MaxQuant. **(b)** PCA plot of TMT-Integrator median-centered gene report from tumor and normal samples. **(c)** Boxplots showing the R-squared value distributions for protein abundance correlation between replicate runs of the NCI and QC samples. FP represents FragPipe, MQ represents MaxQuant, RefRatio represents ratio-to-reference normalization with a real reference, non-norm represents absolute intensity data without ratio-to-reference normalization and MD represents the use of median-centering normalization. **(d)** Violin plots illustrating gene-wise and sample-wise protein-RNA correlations from OmicsEV results across different methods, with median correlation coefficients labeled for each method. VirtualRatio represents ratio-to-reference normalization with a virtual reference. **(e)** Boxplots comparing the noise to signal ratio (NSR) levels between protein and peptide levels in both NCI and QC channels for TMT-Integrator reports. abundance correlation between replicate runs of the same QC sample

In terms of protein-RNA correlation evaluation, FragPipe under all settings, including default settings for datasets with multiple plexes and common bridging sample (‘FP RefRatio MD’) demonstrated good correlation between protein and RNA data, both sample-wise (protein-RNA correlation within the same sample) and gene-wise (protein-RNA correlation across all samples), as shown by the median correlation coefficients in **Table 1** and the overall distribution for selected settings as displayed in **Figure 3d**. Of note, median centering significantly improved gene-wise correlation, from 0.28 to 0.41, in FragPipe (note that sample-wise correlation is not affected by the normalization choice). Given that observation, and because MaxQuant does not perform additional normalization, we applied the median centering to MaxQuant results ourselves (see **Methods**). MaxQuant, with normalization to reference and median centering (‘MQ RefRatio MD’), has shown the same gene-wise protein-RNA correlation of 0.41 as FragPipe under similar settings (‘FP RefRatio MD’). However, the importance of estimating absolute protein abundances (as compared to just using ratios-to-reference as the final output) becomes evident from inspecting the sample-wise correlation between RNA and protein data. Unlike gene-wise (i.e. intra sample) correlation that can be calculated using normalized values such as ratios, comparing RNA and protein abundance within the same sample requires absolute protein abundance estimation. Indeed, as shown in **Table 1** and **Figure 3d**, TMT-Integrator computed abundances (obtained by converting the ratios back to absolute intensities, see **Methods**) shows a notably higher mean gene-wise abundance correlation, R_sample-wise_, compared to that using MaxQuant results: 0.46 (‘FP RefRatio MD’) vs 0.19 (‘MQ RefRatio MD’). The ability of FragPipe with TMT-Integrator to report absolute protein abundances enables additional biological analyses. For example, as we described in our original ccRCC publication [21], we observed higher sample-wise protein-RNA correlation in tumors associated with certain clinical features such as high tumor grade or mutation status of key genes/proteins, and were then able to link high sample-wise RNA-protein correlation to increased protein translation. Running MaxQuant without normalization to reference (‘MQ NoNorm MD’), which produces output tables with absolute intensities, does increase the sample-wise protein-RNA correlation to 0.41 (albeit still lower than that for FragPipe with TMT-Integrator). However, it comes at a cost of a significant decrease in gene-wise protein-RNA correlation, to 0.21 (**Table 1**, **Figure 3d**).

We also used OmicsEV to evaluate the results with respect to correlation within (intracomplex) and between (intercomplex) protein complexes (based on CORUM [66]) using the Kolmogorov-Smirnov (KS) test metric. The resulting ‘complex K-S’ score (**Table 1**) measures the ability of different processed datasets to capture biologically relevant information. The rationale behind this evaluation is that the median correlation of abundances of protein pairs that are part of different protein complexes is likely to be close to zero while the median correlation of protein pairs within protein complexes is likely to be higher than zero. Thus, the K-S test statistic can be used to measure the difference between the two types of correlation values. The higher K-S value indicates that the abundance correlation is higher (i.e., more consistent) within protein complexes and lower (i.e., more dissimilar) between protein complexes. FragPipe/TMT-Integrator and MaxQuant both performed best with the normalization to reference sample approach and with MD normalization.

OmicsEV also computes a KEGG pathway membership prediction score (func_auc score). This method evaluates the biological signal in a data table by constructing co-expression networks from each data table and assessing functional category predictions. For a chosen functional category (in this case, from KEGG [67]), proteins/genes annotated to the category form the positive set, while others form the negative set. Subsets of these proteins/genes are used as seeds for random walks through the network to calculate scores for the remaining proteins/genes, where higher scores indicate a closer relationship to the seed proteins/genes. **Table 1** shows the func_auc scores as metrics of prediction performance (the higher the better). FragPipe with TMT-Integrator, using normalization to reference, shows a stronger performance (a functional prediction score of 0.82 for ‘FP RefRatio MD’) compared to that of MaxQuant (a score of 0.72 for ‘MQ RefRatio MD’). This suggests that FragPipe generated protein table may better reflect protein function than that of MaxQuant. **Supplementary** Figures 2c and 2d further support this finding, showing that FragPipe has more KEGG categories (102) where the protein table exhibits more biological signals than the RNA table, compared to MaxQuant’s 63 categories.

As bridging samples may not be included in each plex due to experimental design constraints, we have implemented an option in TMT-Integrator to artificially generate a virtual reference channel, including using the median abundance across the TMT channels in an experiment (see **Methods**). Using the virtual reference channel approach (‘FP VirtualRef MD’) resulted in a similar performance across all metrics as when using the real pooled reference sample. This is an encouraging observation, suggesting the virtual reference approach of TMT-Integrator can be used to process and harmonize data from different cohorts. MaxQuant implements a somewhat similar option (‘MQ VirtualRef’), however, it did not perform as well in our tests (**Table 1**). We also explored the use of an alternative, “weighted ratio” approach for aggregation of PSM-level ratios-to-reference to the gene/protein level. The weighted approach in TMT-Integrator in FragPipe resulted in slightly worse gene-wise correlation between protein and RNA data compared to the conventional median ratio aggregation (0.38 vs 0.41), and less clean separation between tumor and normal samples in the PCA plots (see **Supplementary File 1**). Furthermore, we observed somewhat higher noise-to-signal (NSR), computed as the standard deviation ratios between the QC samples (the five NCI-7 and eight QC) and patient samples. Surprisingly, in MaxQuant, the weighted ratio approach showed more noticeably worse results across all metrics (**Table 1**; **Supplementary File 1**).

The analysis described above is based on the use of data summarized from PSM to the gene/protein level. Such a summarization has an advantage in that inaccurately quantified peptides are removed from the analysis as outliers during the aggregation from PSM to protein. As a drawback, it inevitably results in the loss of biological information, e.g., due to inability to quantify different protein products of the same gene (e.g. different splice isoforms)[68, 69]. Thus, TMT-integrator, in addition to the summarization to the gene/protein level, also creates peptide-level summary reports. We did note, however, higher NSR values for the NCI-7 and QC technical replicates when calculated using the abundances at the peptide compared to protein levels (**Figure 3e**).

### Evaluation of TMT-Integrator using a spiked-in benchmark dataset

A publicly available *E*. *coli* dataset with 13 spike-in proteins was downloaded from ProteomeXchange (identifier: PXD005486), which included 24 fractions acquired with MS2-level quantification and 24 fractions acquired with MS3-level quantification. The analyses of both MS2 and MS3 data were conducted separately using FragPipe/TMT-Integrator and compared to MaxQuant and Proteome Discoverer. As shown in Table 2, each of the three tools detected all 13 spiked-in proteins in both the MS2 and MS3 data, while TMT-Integrator detected the most *E. coli* proteins.

**Table 2.**
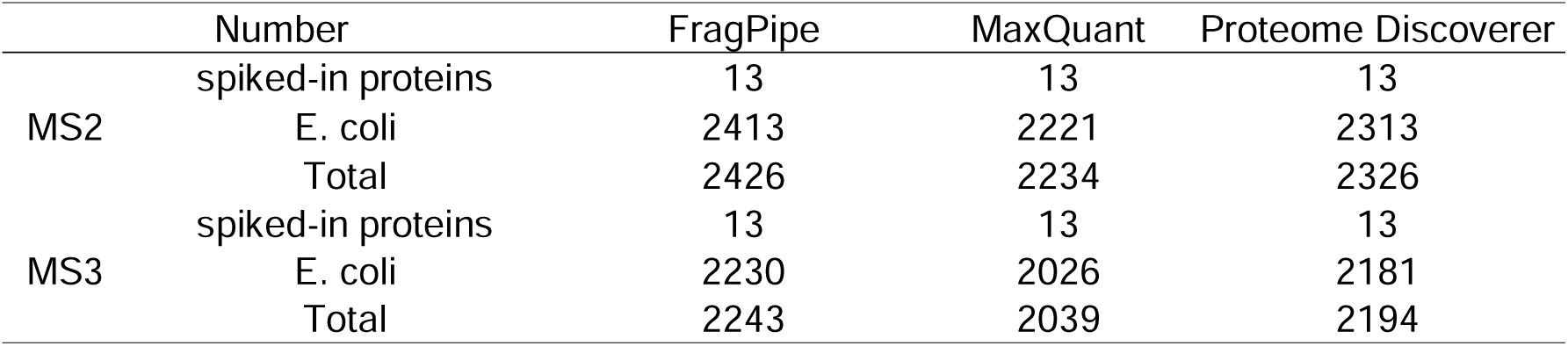
The number of genes in the MS2 and MS3 datasets reported by FragPipe, MaxQuant and Proteome Discoverer.

First, we used the abundances of all *E. coli* proteins to investigate whether median-centering ratio normalization would improve quantitative accuracy. Given that the *E. coli* sample amount across the ten channels is equal, the expected protein ratios for all *E. Coli* proteins should be 1. We therefore calculated the protein ratios between any two of the ten channels (See **Methods** for detail). Comparing the ratio distributions of MS2 and MS3 data (with and without median centering) in **Figure 4a**, we observed that FragPipe and MaxQuant have similar ratio distributions, and the median-centered ratios were much closer to 1, indicating the importance of ratio normalization prior to downstream statistical analysis. Therefore, the median-centered abundances will be used in subsequent analyses unless otherwise noted. It is also worth noting that the ratios reported by Proteome Discoverer were close to 1 regardless of whether median centering was used because peptide-level normalization was applied within Proteome Discoverer.

**Figure 4.**
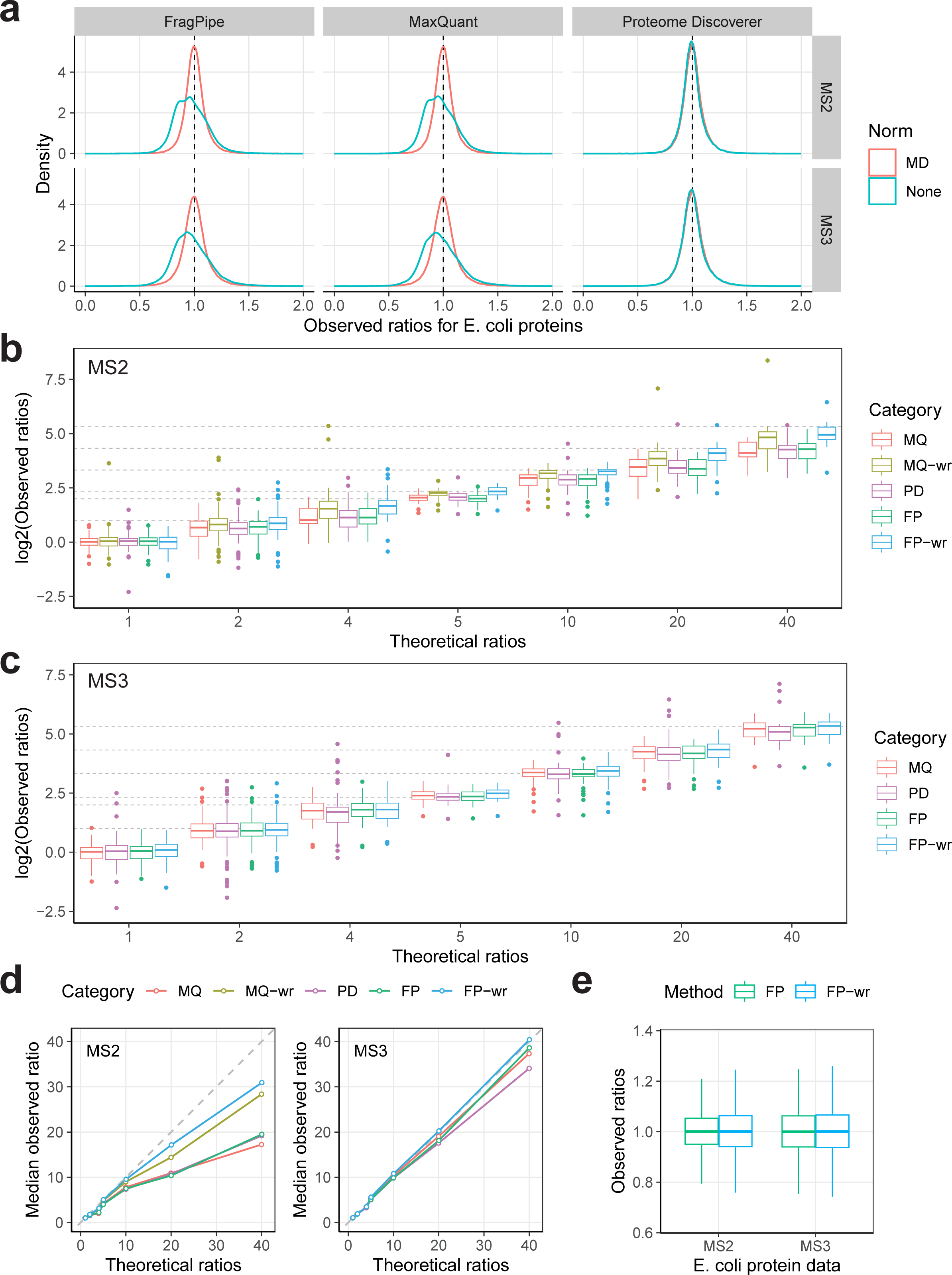
Performance evaluations on the spiked-in dataset. **(a)** Impact of median-centering normalization on E. coli protein quantification accuracy. Density plots show the observed ratio distribution of E. coli proteins across FragPipe, MaxQuant, and Proteome Discoverer using MS2 and MS3 data. Line colors indicate whether the median-centering normalization was used. **(b)** Evaluation of protein quantification accuracy using the 12 spiked-in proteins. Boxplots show the observed ratio distribution of 12 spiked-in proteins compared to theoretical ratios (grey dashed lines) in MS2 data across different methods. Box colors represent the respective methods, with MQ stands for MaxQuant, MQ-w for the weighted method in MaxQuant, PD for Proteome Discovery, FP for FragPipe, and FP-w for the weighted method in FragPipe. **(c)** Same as **(b)** for MS3 data. **(d)** Line charts showing the agreement of median observed ratios with theoretical ratios. Each dot represents a median value of observed ratios. **(e)** Comparison of the observed ratio distributions for E. coli proteins between FragPipe’s conventional median ratio method and weighted ratio method.

Next, we used the abundances of the 13 spiked-in proteins to evaluate performance across the three tools on MS2 and MS3 data. Detailed spiked-in amounts for the 13 proteins in each channel are described in [45]. In brief, the 13 proteins were spiked into the E. coli samples at different amounts to represent fold changes of 1, 2, 4, 5, 10, 20 and 40. Due to varying amounts of Albumin, we excluded it and used the other 12 proteins for performance evaluation. Since the 13 proteins were injected in known amounts, the reported protein abundance ratios should reflect the amounts added, with better agreement between theoretical and observed protein ratios indicating higher accuracy. We calculated the spiked-in protein ratios between any two of the ten channels (**Supplementary Table 1)**. **Figures 4b and 4c** show the observed and theoretical ratios of the seven groups in MS2 and MS3 data, respectively (the evaluation results without median centering are provided in **Supplementary** Figure 3a). On MS2 data, all investigated tools showed noticeable deviations from the expected ratios for all groups except the group representing a fold change of 1. The ratio suppression effect is more evident in **Figure 4d**, which plots the median observed ratios for the seven groups. Notably, in MS2 data, the weighted method in FragPipe consistently outperformed that in MaxQuant across all groups, especially for fold changes of 20 and 40. The deviation became more pronounced in groups representing larger fold changes, such as 20 and 40, indicating all tools suffered from a ratio compression to some extent. We also observed that using the weighted median approach performed better for accurate quantification in MS2 data, with the observed ratios being much closer to the expected ratios, compared to the conventional median approach. In contrast, the advantage of weighted approach over conventional median approach on MS3 data was far less noticeable. In MS3 data, all methods showed overall good agreement with the expected ratios except one group, ratio 4, which may be due to inaccurate estimation of the amount of spiked-in proteins. Note that we excluded the MaxQuant weighted method in MS3 data due to poor performance (see **Supplementary** Figure 3b). Among all tools, Proteome Discoverer appeared to be less accurate with MS3 data, showing slightly larger deviations and more outliers compared to MaxQuant and FragPipe. The increased number of outliers in Proteome Discoverer (as clearly visible in **Figure 4c**) is likely due to its summed intensity approach (summing the reporter ion intensities from all PSMs to their corresponding protein).

We further investigated the difference between the conventional median and weighted approach performance using the E. coli proteins and proteins from the ccRCC proteome data. **Figure 4e** shows that the ratio distributions for E. coli proteins in MS2 and MS3 data are similar between the conventional median and weighted approaches in FragPipe/TMT-Integrator, all centered around the expected ratio of 1. However, the conventional median approach exhibited less deviation, with the interquartile range (IQR) in the boxplots consistently showing narrower distributions compared to the weighted approach for both MS2 and MS3 data. This can also be seen in the density plot (**Supplementary** Figure 3c), where the conventional approach distribution is higher and narrower than the weighted approach, a similar trend also evidenced in MaxQuant results. For the ccRCC proteome data, we examined the differences in protein NSR values when comparing the conventional median and weighted approaches. The conventional median approach clearly performed better, resulting in more proteins with smaller NSR values (**Supplementary** Figure 3d). Based on these results, we conclude that the weighted approach may be more suitable for MS2 data and when it is important to accurately quantify proteins showing very large fold changes (e.g., in affinity purification - MS experiments comparing bait protein purification against negative controls). At the same time, the conventional median approach is likely to be more suitable for common global proteome profiling experiments and, therefore, selected as the default option in TMT-Integrator.

### Evaluation of the phosphorylation site reports

Besides generating gene, protein and peptide reports, TMT-Integrator also creates site-level reports for specified modifications in PTM-enriched proteomic data, which have been used in multiple studies, e.g. [21, 24, 48]. In this process, TMT-Integrator uses PTM localization probabilities from PTMProphet in FragPipe and summarizes quantification information at the level of multiple and single modification sites. The details on how the multi-site and single-site reports are generated, along with illustrations of multi-site and single-site indexes, are described in the **Methods** section and **Supplementary Figure S1**.

To evaluate the performance of FragPipe with TMT-Integrator in analyzing PTM-enriched data, we used the ccRCC phosphorylation-enriched dataset. As the whole proteome dataset discussed earlier, its phosphorylation-enriched counterpart contains tumor and normal samples, alongside eight NCI and five QC samples. **Figure 5a** shows the number of identifications of phosphorylated peptides and proteins at different levels. A total of 9045 phosphorylated proteins corresponding to 9037 genes were identified, with 4148 proteins and 4145 genes detected in all samples. At the peptide (sequence) level, 48220 phosphorylated peptides were identified, of which 9389 were quantified across all samples and 24070 were quantified in more than 50% samples. At the peptidoforms level (considering all reported site configurations and counting localized and unlocalized configurations as separate entries), there were 108310 phosphorylated peptidoforms (multi-site report) and 60733 localized sites (single-site report), of which 17292 sites were detected in at least 50% of the samples. As expected, peptide level data exhibited better data completeness compared to multi-site and single-site data, reflecting the fact that in many instances a phosphopeptide could be consistently identified across many samples but sites of phosphorylation confidently localized in a subset of those. This also explains why some past studies, e.g. [21], chose to perform certain downstream analyses using phosphopeptide and not site-level data. However, most downstream analyses of PTM-centric data require a single-site level quantification table. For example, PTM-SEA [63] uses quantification information from specific sites to check for enriched sites in pathways and infer kinase activity. Thus, we decided to conduct performance evaluation and comparison with MaxQuant using the single-site reports.

**Figure 5.**
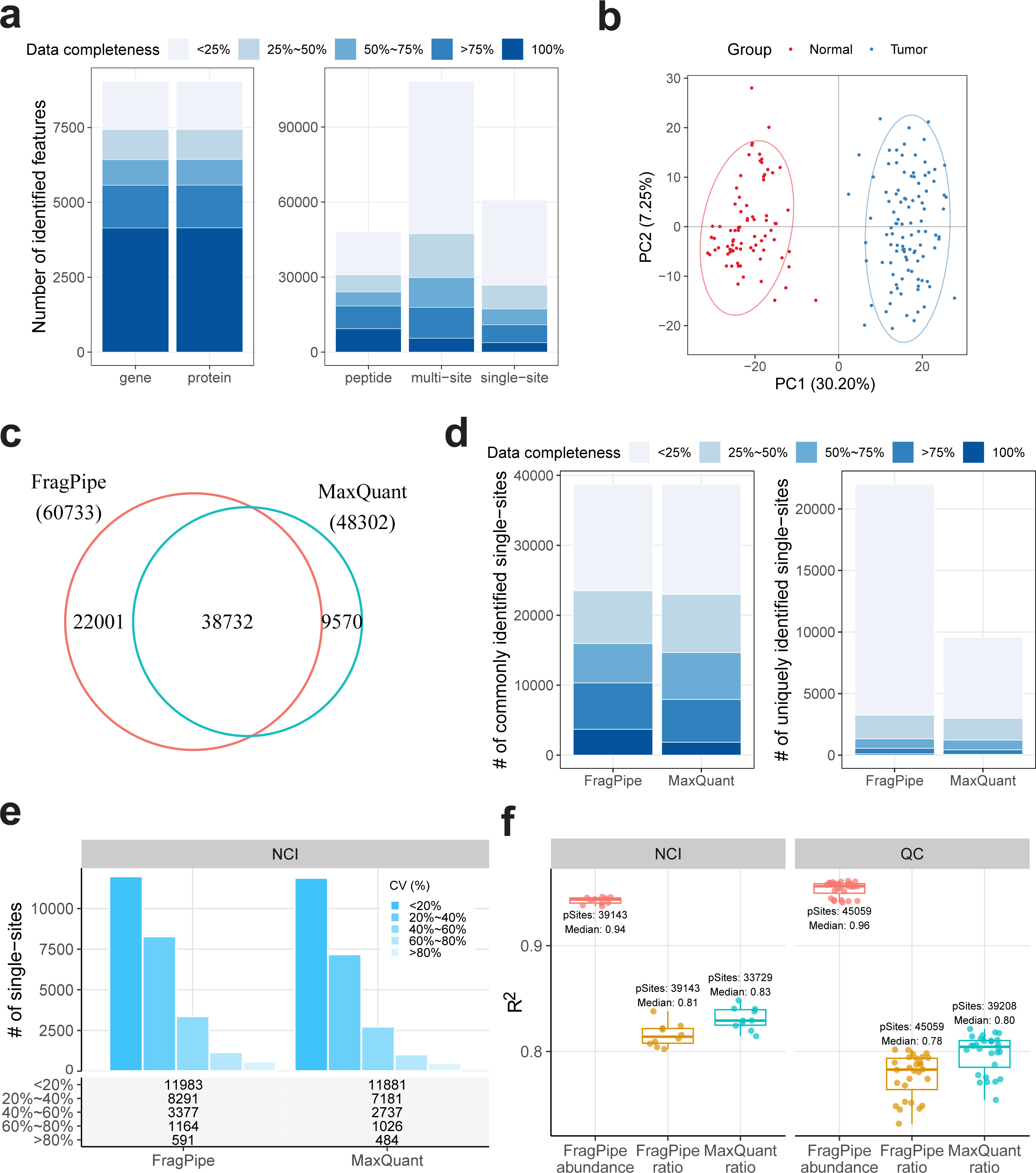
Performance evaluations on the ccRCC phosphorylation-enriched dataset. **(a)** Identification performance across all report levels and all samples. The number of IDs with varying data completeness is represented by different color shades, with darker blue indicating higher completeness. IDs detected in all samples are marked as 100%, and IDs detected in less than 25% of samples are marked as <25%, with the similar logic applied to other groups. **(b)** PCA plot of TMT-Integrator median-centered single-site data from tumor and normal samples. **(c)** The number of single-sites identified by FragPipe and MaxQuant. **(d)** Data completeness comparison between FragPipe and MaxQuant for commonly and uniquely identified single-sites. **(e)** Comparison of single-site variation (CV) distributions in NCI channels between FragPipe and MaxQuant, with bars representing the number of single-sites in each CV group. **(f)** Evaluation of quantification consistency across FragPipe abundance and ratio single-site reports and MaxQuant single-site ratio data in NCI and QC channels. Boxplots show the distribution of R-squared values from linear model fitting of single-site quantifications for each sample pair in NCI and QC channels. The labeled text shows the total number of single-sites and the median R-squared value.

We first performed PCA on the data from tumor and normal samples. The single-site PCA plot (**Figure 5b**) shows that samples clustered into two distinct groups along PC1, clearly separating tumor and normal samples as expected, indicating strong differences in site abundance between the two groups. Consistently, PCA plots for other levels (**Supplementary** Figure 4a) show a highly similar separation, suggesting that TMT-Integrator reports provide reliable quantification at all levels. Since MaxQuant does not directly generate single-site reports. We applied the same process as in TMT-Integrator to generate a single-site report from MaxQuant Phospho (STY) Sites.txt (the script is provided with this study, **Code Availability**). The generated MaxQuant single-site data was median-centered, and the PCA plot **(Supplementary** Figure 4b) showed a similar separation between tumor and normal samples. **Figure 5c** shows the overlap of phosphorylation sites identified by FragPipe and MaxQuant. 38732 phosphorylation sites were detected by both software tools, and 22001 and 9670 sites were uniquely detected by FragPipe and MaxQuant, respectively. As expected, uniquely identified sites were mostly detected in only a few samples, likely reflecting their lower abundance. We then compared the data completeness of both commonly identified sites and uniquely identified sites (**Figure 5d**). For the commonly identified sites, FragPipe resulted in a larger number of fully detected sites across all samples than MaxQuant, suggesting that FragPipe performs better in consistently detecting PTM sites across the cohort, in agreement with a previous report [70].

Next, we evaluated the consistency and robustness of phosphorylation site quantification using replicates of the NCI and QC samples. We computed the coefficient of variation and the ratio correlation among the NCI and QC samples. **Figure 5e** and **Supplementary** Figure 4c show the number of phosphorylation sites in different CV groups for NCI and QC samples, respectively. FragPipe consistently quantified more sites in each CV group than MaxQuant, while both showed similar distributions, indicating comparable performance. The ratio correlation was assessed by fitting a linear regression to each pair of samples and evaluating the fit using R-squared (R²). An essential feature of TMT-Integrator is that it produces both ratio reports and absolute abundance reports and the difference in their data scales can result in slightly different linear fitting results. Thus, we included both abundance and ratio reports for R² calculation and compared them with MaxQuant ratio data. **Figure 5f** shows that both FragPipe abundance and ratio reports had good quantification consistency, with FragPipe computed abundances getting the highest R² and FragPipe computed ratios having R^2^ values similar to MaxQuant. **Supplementary** Figure 4d illustrates the linear fittings for each pair of samples and includes extra Pearson correlation tests, further demonstrating the high consistency of site-level quantification in FragPipe. It is worth noting that both ratio and abundance reports have the same data variation, and the choice will not affect PCA, CV or differential analysis results. However, when the goal of the analysis is to quantitatively integrate PTM site data with proteomic or transcriptomic data (which are in absolute abundance scale), the use of abundance reports (instead of ratios) is recommended. Overall, we observed that FragPipe with TMT-Integrator identifies more phosphorylation sites than MaxQuant while offering comparable site quantification performance.

### Evaluation of Astral and TMT 35-plex datasets

Advancements in mass spectrometry instrumentation and the multiplexing capability of isobaric labeling have progressed significantly in recent years. We utilized two datasets representing these latest advances to showcase the versatility of FragPipe/TMT-Integrator in analyzing data from cutting-edge instruments and novel isobaric labeling reagents. The first dataset comprises 18 cell lysate samples collected using an Orbitrap Astral mass spectrometer in both DDA and DIA modes, facilitating a direct comparison between TMT and DIA quantification using the Astral instrument. The second dataset includes 32 cell lysate samples labeled with TMT 35-plex reagents. Both datasets allow for evaluating the accuracy and consistency of quantification using TMT-Integrator.

The Astral datasets from Liu et al. [49] were employed to assess FragPipe’s performance in quantifying Astral data. Both TMT and DIA achieved deep proteome coverage, with 10,353 proteins (counting the number of proteins at the gene level) quantified in the DIA data, 10,031 proteins quantified in the TMT data, and 9,522 proteins quantified by both methods. In addition to identifying 322 more proteins, the TMT data exhibited fewer missing values across all samples. As shown in **Figure 6a**, TMT quantified 10,024 proteins with no missing values and another seven proteins in more than 50% of the samples. In contrast, DIA quantified 8,169 proteins (78.9% of all proteins) with no missing values, 1,689 in more than 50% of the samples, and 495 in less than 50% of the samples.

**Figure 6.**
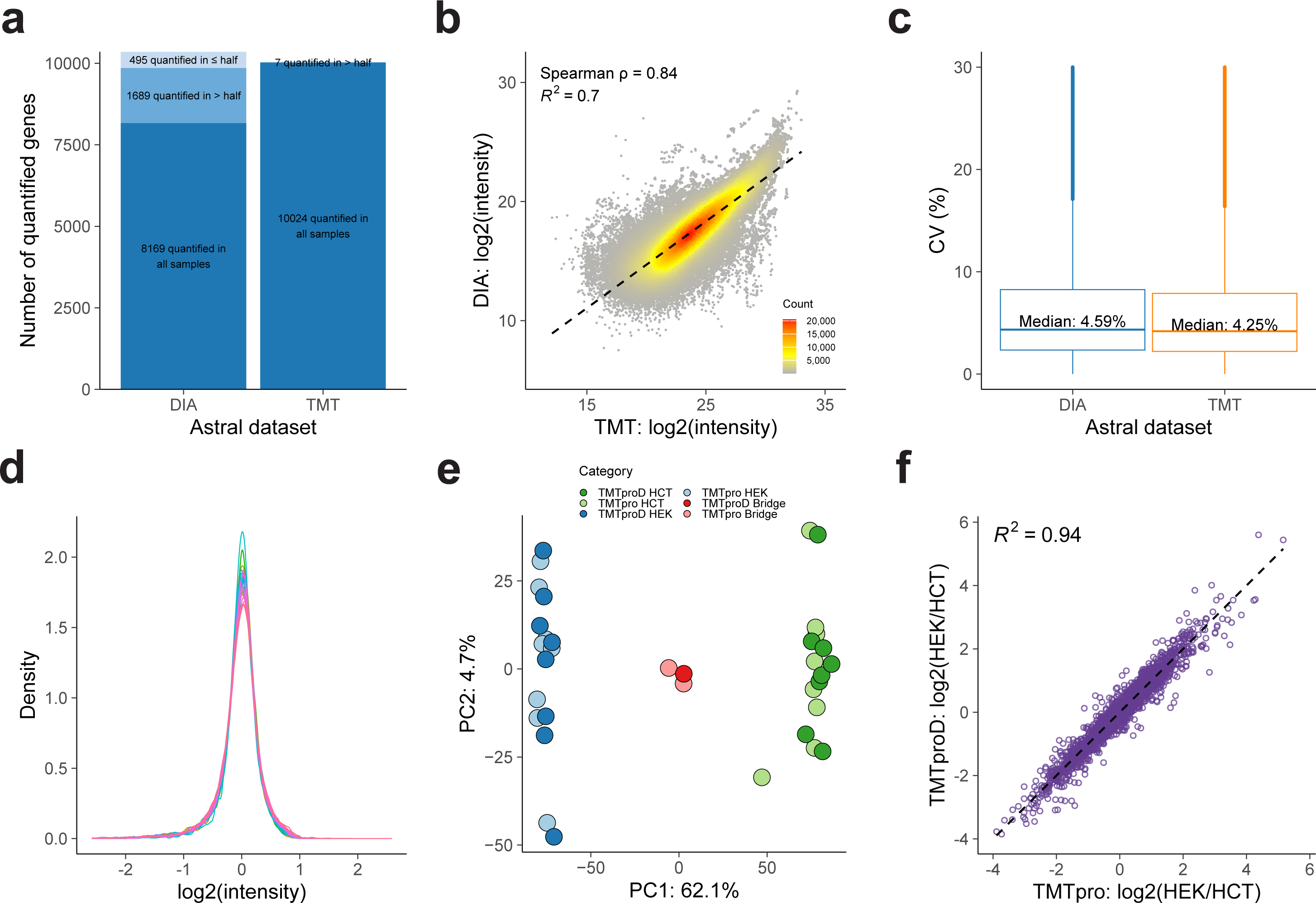
Performance evaluations on the Astral and TMT 35-plex datasets. **(a)-(c)** for Astral datasets and **(d)-(f)** for TMT 35-plex dataset. **(a)** Bar plot showing the number of quantified genes in Astral TMT and DIA datasets across all 18 samples. Proteins are divided into three categories: those quantified in all samples, in more than half of samples, and in less than half of samples. The number of proteins in each category is represented by a different color shade. **(b)** Scatter plot showing the correlation between TMT and DIA measurements for each protein across samples. Each dot represents a protein in a specific sample and is color-coded based on the number of overlapping dots. The black dashed line indicates the overall linear fit. **(c)** Boxplots comparing the combined CV distributions for TMT and DIA, calculated from triplicates for each cell line. The box in each plot captures the IQR with the bottom and top edges representing the Q1 and Q3, respectively. The median (Q2) is indicated by a horizontal line within the box. The whiskers extend to the minima and maxima within 1.5 times the IQR below Q1 or above Q3. **(d)** Density plots displaying the intensity distribution of all samples labeled with both deuterated and non-deuterated reagents, using the raito_gene_MD.tsv report. **(e)** PCA plot presenting sample clustering according to cell type and labeling reagent. TMTproD represents sample labeling with deuterated reagents and TMTpro represents non-deuterated reagents. The common reference samples are denoted as Bridge. **(f)** Scatter plot illustrating the correlation of log2 ratios of HEK to HCT between deuterated and non-deuterated labeling types.

We then assessed the concordance of protein abundance between TMT and DIA measurements for proteins quantified by both methods. In **Figure 6b**, we compared the log2 intensities of TMT and DIA measurements for each protein across all samples. Each dot represents a protein in a specific sample, color-coded by the number of overlapping dots. The overall agreement between TMT and DIA measurements is indicated by a Spearman correlation coefficient of 0.84 and a linear fit with an R-squared value of 0.7, demonstrating a strong correlation.

We also compared the distribution of coefficients of variance (CV) between TMT and DIA measurements. CVs were calculated from triplicates of each condition (cell line and treatment), and the combined CV distributions from all cell lines are shown in **Figure 6c**. The median CV in DIA is slightly higher than in TMT (4.59% vs. 4.25%). Overall, we demonstrate that FragPipe/TMT-Integrator achieves deep proteome coverage and precise quantification by leveraging the fast acquisition rate of the Astral mass spectrometer. Additionally, the FragPipe platform enables consistent analysis of data using both quantification strategies, TMT and DIA.

The TMT 35-plex dataset from Zuniga et al. [58] was used to illustrate how TMT-Integrator can handle higher-order multiplexing data labeled with both deuterated and non-deuterated TMTpro reagents, allowing simultaneous analysis of up to 32 samples. The original study noted that using deuterium isotopes may cause a slight retention time shift in co-eluted peptides, affecting reporter ion quantification accuracy and leading to batch effects. They suggested using three bridge channels - two in the non-deuterium subplex and one in the deuterium subplex - to normalize and remove these effects. We proposed that TMT-Integrator’s virtual reference approach can achieve similar results.

We started by comparing intensity distributions across deuterated and non-deuterated channels. **Figure 6d** shows bell-shaped curves for all channels, indicating similar intensity distributions for both labeling types with no noticeable distortions. We further examined this using PCA analysis. **Figure 6e** reveals clear separation of HCT and HEK cell lines along PC1, with no distinction between samples labeled with non-deuterated and deuterated reagents. Additionally, three bridge samples from both labeling reagents cluster closely.

Together, these results demonstrate that TMT-Integrator ensures accurate quantification without batch effects between deuterated and non-deuterated labeling.

We also assessed the consistency of quantification between labeling types by examining the fold change in protein abundance between HEK and HCT cell lines. We compared the log2 ratios of HEK to HCT and fitted a regression line to assess their agreement. The scatter plot in **Figure 6f** shows a strong correlation (R-squared value of 0.94), indicating consistent quantification between labeling types. Our results with the TMT 35-plex data demonstrate that TMT-Integrator provides accurate and consistent quantification. Notably, the virtual reference approach in TMT-Integrator does not require bridge samples, increasing the potential number of biological samples profiled within each plex. A more in-depth investigation of normalization methods for TMT 35-plex data and the potential benefits of using bridge samples will be explored in future work as more data becomes available.

## CONCLUSION

In this study, we provided a comprehensive evaluation of FragPipe with TMT-Integrator as a robust computational platform for isobaric quantitative proteomics using TMT and iTRAQ. Through analyses of both large-scale clinical datasets, such as the clear cell renal cell carcinoma (ccRCC) dataset, and controlled experimental datasets, such as the E. coli spike-in tests, we demonstrated that FragPipe with TMT-Integrator offers several advantages over existing platforms like MaxQuant and Proteome Discoverer. Our findings illustrate that FragPipe consistently delivers high coverage and accurate quantification across diverse sample cohorts, effectively handling the complexities inherent in multiplexed proteomic experiments. Moreover, our analyses of both Astral and TMT 35-plex datasets demonstrate that FragPipe with TMT-Integrator is versatile, flexibly supporting a wide range of isobaric labeling data from state-of-the-art instruments to the latest multiplexing reagents, consistently providing precise and accurate quantification.

The inclusion of features like reference and virtual reference normalization approaches ensures its applicability to a broad spectrum of experimental designs, which facilitates data harmonization across different cohorts. Additionally, FragPipe’s capability to derive absolute protein abundances from ratio data provides valuable insights, enabling more comprehensive biological interpretations that are not as feasible with purely ratio-based quantifications. The evaluation of phosphorylation data further underscores FragPipe’s utility, revealing improved identification and quantification of post-translationally modified sites compared to other tools.

Future studies can refine the platform’s abilities in processing higher multiplex (35-plex) [58] data and further enhance site-level report generation by incorporating across-sample site localization analysis. This entire pipeline can be conveniently executed using the graphical user interface of FragPipe (http://fragpipe.nesvilab.org/) or via the command line for high-throughput applications. Moreover, the output from TMT-Integrator is compatible with downstream tools such as FragPipe-Analyst [71] and MSstats [35], offering extensive flexibility for subsequent data analysis.

## METHODS

### Datasets

*Clear cell renal cell carcinoma (ccRCC) whole proteome and phosphoproteome datasets.* The ccRCC proteome and phosphoproteome datasets are available from the CPTAC Data Portal at https://cptac-data-portal.georgetown.edu/study-summary/S044. A total of 110 ccRCC tumor samples, 84 paired normal adjacent tissue (NAT) samples, 8 aliquots of a non-CPTAC kidney tumor sample (labelled ‘QC’), and 5 aliquots of the NCI-7 mix of cell lines (samples labelled ’NCI7’) were randomly assigned to the 23 TMT 10-plex sets in the proteome and phosphoproteome datasets. The last channel in each TMT 10-plex was reserved for the bridging (reference) sample, created from the pool of all patient samples in the cohort. Each experiment in the proteome dataset has 25 fractions, while each experiment in the phosphoproteome dataset has 13 fractions, obtained using offline LC fractionation. As described in the original publication [21] and [48], 7 tumor samples were later determined to be of non-ccRCC subtype, and removed from the final analysis, along with the QC and NCI samples added for quality control. The evaluation of the results was based on the set of 103 ccRCC tumors and 84 NAT samples and, where noted, additional analyses were performed using the QC and NCI7 samples.

*E. coli spike-in dataset.* The *E. coli* dataset is composed of ten whole cell lysate samples with 12 recombinant human proteins plus bovine serum albumin spiked in to generate fold changes of 1, 2, 4, 20, and 40 among the TMT 10-plex labeled samples. Reporter ion data acquisition was performed at both MS2 and MS3 levels, resulting in two sets of data, each containing 12 fractions. The dataset was downloaded from the PRIDE repository with identifier PXD005486.

*Astral datasets.* The Astral datasets include 18 human cell lysate samples from four cell lines (IHCF, HCT116, HeLa, and MCF7). The IHCF cell line was additionally treated with two concentrations of H2O2, resulting in six distinct conditions, each in biological triplicate. These samples were analyzed using a Thermo Fisher Scientific Orbitrap Astral mass spectrometer with both TMT and DIA strategies. In the TMT experiment, samples were labeled with TMTpro18-plex reagents, and data was collected after fractionation into 24 peptide fractions. For the DIA experiment, samples were analyzed using a 60-minute gradient, and data was collected with and without FAIMS. Only the data without FAIMS was used for performance evaluation in this study, as it yielded better results as described in the original publication [49]. The datasets were downloaded from the PRIDE repository with identifier PXD058918.

*TMT 35-plex dataset.* The TMT 35-plex dataset consists of 35 human cell lysate samples, of which 18 are labeled with non-deuterated TMTpro reagents and 17 with deuterated TMTpro reagents (TMTproD). For each type of reagent, there are eight replicates of the HEK116 and HCT293T cell lines. Besides, there are three common references (bridge sample) derived from equal amounts of pooled cell lysates across all replicates from both cell lines: two in non-deuterated labeling and one in deuterated labeling. Although labeled with different reagents, all samples were analyzed together in a single experiment using a Thermo Fisher Scientific Orbitrap Exploris 480 mass spectrometer, and data was collected in 12 factions. The dataset was downloaded from the PRIDE repository with identifier PXD054559.

### FragPipe data processing

*ccRCC whole proteome and phosphopeptide-enriched datasets*. All MS files in mzML format were processed via FragPipe v. 22, a newer version than that used in the previous publication. Unless specified, default FragPipe parameters were applied (‘TMT10-bridge’ or ‘TMT10-phospho-bridge’ workflows). MSFragger v. 4.1 was used to search the spectra against the reviewed human protein sequences downloaded from UniProt (retrieved 2024-08-09). Decoy and contaminant protein sequences were added using the Philosopher toolkit via FragPipe. Searches were performed with oxidation of methionine and protein N-terminal acetylation as variable modifications and cysteine carbamidomethylation as a fixed modification. Restrict trypsin was selected as the enzyme, allowing up to two missed cleavages, and the peptide mass was limited to a maximum of 5000 Da. The precursor mass tolerance was set to 20 ppm, C12/C13 isotope errors allowed (−1/0/1/2/3). Mass calibration and parameter optimization [39], and MS/MS spectral deisotoping [50] were enabled. Cysteine carbamidomethylation (+57.0215) and lysine TMT labeling (+229.1629), were specified as fixed modifications. Methionine oxidation (+15.9949), and peptide N-terminal TMT labeling, and serine TMT labeling were specified as variable modifications. For phosphopeptide enriched data, the set of variable modifications also included phosphorylation (+79.9663) of serine, threonine, and tyrosine residues, but excluded the serine TMT labeling, and with C12/C13 isotope errors parameter set to (0/1/2). MSFragger search results were processed using MSBooster [51] (except phosphoproteomics data), Percolator [52], PTMProphet (phosphopeptide-enriched data) [53], and ProteinProphet [54]. Resulting PSM lists for each TMT plex were filtered using 1% PSM-level and global 1% protein-level FDR using Philosopher [40]. Precursor ion MS1 intensity and TMT reporter ion intensities were extracted from the MS and MS/MS scans, respectively, using IonQuant [41]. The precursor ion purity scores were calculated using the intensity of the sequenced precursor ion and that of other interfering ions observed in MS1 data (within a 0.7 Da isolation window). All supporting information for each PSM, including the accession numbers and names of the protein/gene selected based on the protein inference approach with razor peptide assignment and quantification information (MS1 precursor-ion intensity and the TMT reporter ion intensities), were summarized in the output PSM.tsv files, one file for each of the 23 TMT plexes. Both ccRCC global proteome and phosphoproteome datasets contain the pooled sample as one of the channels in each plex, and the pooled samples were specified as the reference samples (default). FragPipe was also run using other options, including the virtual reference approach (i.e. without specifying a reference sample) as noted in the main text.

*E. coli spike-in dataset*. Spectral files from 24 fractions of the *E. coli* data (12 from the MS2 dataset and 12 from MS3) were converted from Thermo RAW to mzML file format using ProteoWizard [55]. The protein sequences of *E. coli* K12 were downloaded from UniProt [56] (accession number UP000000625), and contaminants were added along with the reversed protein sequences using Philosopher. The 13 spiked-in proteins were manually added to the *E. coli* database along with their reversed protein sequences, generating a total of 8828 sequences (target and decoy). The converted mzML files were searched against the *E. coli* database using FragPipe v. 22 as described above for the ccRCC whole proteome dataset, except in MSFragger TMT-labeling of peptide N-termini was set as fixed modifications. TMT reporter ion intensities were extracted from either MS/MS scans (MS2 dataset) or MS3 scans (MS3 dataset) using 0.002 Da extraction window. TMT-Integrator was run using the virtual reference option.

*Astral dataset*. The reviewed human protein sequences were downloaded from UniProt (accession number UP000005640) on 2025-01-26, appended with common contaminants. Decoy protein sequences were added using Philosopher. For the TMT dataset, RAW files (without conversion to mzML) were searched against the Human database using FragPipe (v23.0) with the default TMT18-Astral workflow. Most search parameters stayed the same as those for the ccRCC whole proteome dataset. In MSFragger, the TMT modification mass was set to +304.20715. MSBooster was disabled. In TMT-Integrator, instead of the default intensity-based filter, a minimal resolution of 45000 and minimal single-to-noise (SNR) of 1000 were applied to filter PSMs. The “best PSM” option was disabled, the “min intensity (percent)” was set to 0, and the “mass tolerance” was set to 10 ppm. For the DIA dataset, MS raw files were converted into mzML format as previously described. The mzML files were searched against the same database using FragPipe (v23.0) with the default DIA_SpecLib_Quant workflow. In short, MSFragger-DIA [57] was used to search the DIA data directly, and the search results were processed using MSBooster for deep learning-based score calculation, Percolator for rescoring and posterior error probability calculation, ProteinProphet for protein inference, Philosopher for FDR filtering, and EasyPQP for spectral library building. The spectral library was filtered to 1% global peptide and protein FDR. The resulting library was passed to DIA-NN to extract and quantify precursors, peptides, and proteins from the DIA data.

*TMT 35-plex dataset*. MS raw files were converted into mzML format and analyzed using FragPipe (v23.0) with the same Human database as the Astral datasets. Most search parameters were kept the same as those for the ccRCC whole proteome dataset, except in MSFragger the TMT-labeling modification mass was set to +304.20715 for both non-deuterated and deuterated TMTpro reagents, and an additional new feature in TMT-Integrator (v6.1.0) was applied to achieve accurate quantification from the TMT 35-plex dataset. The original publication [58] noted that the deuterium isotope effect can result in a slight shift in retention time for coeluted peptides, which can impact reporter ion-based quantification and cause batch effects. They also demonstrated that this effect can be effectively mitigated by treating non-deuterated and deuterated channels as two separate subplexes during normalization. To ensure accurate quantification in TMT 35-plex data, we made an adjustment in TMT-Integrator by splitting the 35 channels into non-deuterated and deuterated subplexes before doing anything else. This adjustment allows TMT-Integrator to proceed with two subplexes in a way similarly to a typical multi-plex dataset without any changes to its core algorithm. To generate quantification reports from this dataset using TMT-Integrator, we used the virtual reference approach, where a virtual reference channel was created for each subplex by averaging intensities across channels within that subplex for each PSM. PSM ratio-to-reference normalization was then conducted separately for each subplex using its virtual reference channel, resulting in two sets of ratio data. The rest of the processing follows the steps.

### MaxQuant data processing

Raw MS/MS data were processed using MaxQuant [37, 59] (version 2.6.4.0; released July 28, 2024) with the Andromeda search engine [60] against the same human and *E.coli* protein sequence databases as used in FragPipe (see above). Enzyme specificity was set to “Trypsin/P” and up to two missed cleavages were allowed. Carbamidomethyl (C) was set as a fixed modification, “Oxidation (M)” and “Acetyl (protein N-term)” were set as variable modifications. For the ccRCC phosphoproteome dataset, “Phospho (STY)” was also added as variable modifications. For both ccRCC global proteome and phosphoproteome datasets, the type was set to “Reporter ion MS2” and the isobaric label was set to “10plex TMT”. FDR cutoff of 1% was used at the PSM and protein levels. For the phosphopeptide identification in ccRCC phosphoproteomics dataset, only PSMs passing 1% FDR and with confidently localized phosphosites (localization probability□>□0.75) were used for downstream analysis. For the *E. coli* datasets, the type was set to “Reporter ion MS3” for the MS3 data and “Reporter ion MS2” for the MS2 data. The MaxQuant protein tables of ccRCC global proteome dataset were collapsed to the gene level for the direct comparison with TMT-Integrator gene reports (**Code Availability**). That is, for the proteins corresponding to the same gene, the median abundances were taken. With respect to the normalization approach, all MaxQuant options - “None”, “Ratio to reference channel”, and “Weighted ratio to reference channel” - were tested for the ccRCC global proteome and the *E.coli* datasets. The “Ratio to reference channel” was used for the ccRCC phosphoproteome dataset. Since both ccRCC global proteome and phosphoproteome datasets contain the pooled sample as one of the channels in each plex, we used them as the references when running MaxQuant with “Ratio to reference channel”, and “Weighted ratio to reference” approach. We also applied the virtual reference approaches in MaxQuant to the ccRCC global proteome dataset to compare the results with that from FragPipe virtual reference approach. The *E.coli* dataset does not contain the real reference channel; thus, we applied the virtual reference approach. Other parameters were set to the default values. We also performed an additional, median-centering normalization during the downstream analysis.

### Proteome Discoverer data processing

Proteome Discoverer (PD) v3.0 (Thermo Fisher Scientific) was used for data analysis of the *E. coli* dataset. Since PD adds decoys automatically, MS2 spectra were searched against a target-only subset of the FASTA sequence file used for FragPipe and MaxQuant. The following search parameters were used: MS1 tolerance was set to 10 ppm, and MS2 tolerance was set to 0.02 Da (MS2 dataset) or 0.6 Da (MS3 dataset). Carbamidomethylation of cysteines and TMT labeling of lysine were set as static modifications. Oxidation of methionine, TMT labeling of serine and peptide N-termini, and acetylation of protein N-termini were set as variable modifications. Trypsin was set as the enzyme. Peptides of 7 to 50 residues in length with a maximum of two missed cleavages were allowed. PSMs were subsequently processed using Percolator [52], and identified peptides and proteins were filtered to retain only those that passed ≤1% FDR threshold. Quantification was performed using reporter ion intensities from MS2 or MS3 spectra depending on the dataset. Protein quantification was done allowing unique and razor peptides, with quantification normalized using the total peptide amount option.

### Result Evaluation in OmicsEV

OmicsEV takes reports generated by TMT-Integrator or MaxQuant, along with a sample annotation file, as input and creates a folder that contains datasets to be evaluated. The data quality evaluation components implemented in OmicsEV include missing value distribution, batch effect evaluation, unsupervised clustering analysis, correlation analysis, and phenotype and gene function prediction based on expression data. These analyses produce more than 20 evaluation metrics. All the evaluation results are included in an HTML report, which helps identify the optimal analysis method for the omics dataset under investigation. If the datasets for evaluation were protein expression data, paired mRNA expression data (if available, which was the case here for the ccRCC dataset) could also be used in the evaluation, and vice versa. All these analyses are wrapped into a single function to facilitate easy usage. More details regarding OmicsEV and the source code are available at https://github.com/bzhanglab/OmicsEV. An example script used to analyze the data in this work is provided in the **Code Availability**.

### Noise-to-Signal ratio (NSR) ratio in ccRCC

As described above, the ccRCC datasets included 8 aliquots of a non-CPTAC kidney tumor sample (‘QC’ samples), and 5 aliquots of the NCI-7 cell line mix (‘NCI7’) profiled as part of the sample cohort. These samples were used to evaluate the performance of different pipelines using the coefficient of variance (CV) and the noise-to-signal ratio (NSR) metrics. NSR was calculated for each protein (or peptide, or phosphosite) as the standard deviation across the 5 NCI7 (or 8 QC) samples divided by the standard deviation calculated across the 195 patient samples. Smaller NSR values are indicative of more accurate quantification measurements.

### TMT-Integrator algorithm

TMT-Integrator takes PSM tables generated by Philosopher in FragPipe (**Figure 1a**), one table for each TMT plex, as input and exports an integrated report with columns for sample names and rows for abundances at a user-specified level (including gene, protein, peptide, and modified site), as described below. The index and related information are reported based on the defined levels. The seven steps in TMT-Integrator are explained as follows (**Figure 1b**), concluding with the report generation at various levels (**Figure 1c**):

*Step 1. Filtering.* PSMs with no TMT modification on the peptide, zero intensity in the sample designated as the reference sample (if specified), and precursor ion purity less than the user-specific threshold (50% by default) are first removed for each input table. Then, the reporter intensities across all channels in the plex are summed, and the fraction of PSMs with the lowest summed intensities (by default, lowest 5% for the whole proteome and 2.5% for phosphopeptide-enriched data) are excluded. Since a peptide may have multiple PSMs in the same sample/fraction, only PSMs with the maximum summed intensities are retained (i.e., the best PSMs; default option). PSMs mapping to contaminants are also filtered out. By default, both unique and razor peptides are used for the analysis (controlled by “Peptide-Protein uniqueness” parameter in TMT-Integrator tab of FragPipe). In certain situations, it is useful to restrict the estimation of protein-level expression to unique peptides only. However, restricting the analysis to peptides unique to a single entry in the protein database may lead to a significant loss of peptides, especially in the case of larger databases such as RefSeq or full UniProt (i.e., including unreviewed entries and/or isoforms). Thus, TMT-Integrator also provides a less conservative option (“Peptide-Gene uniqueness”; set to keep all PSMs by default) to remove peptides mapping to a set of proteins representing multiple gene symbols in the protein sequence database. Example of using this option in practice as part a large-scale proteogenomics study can be found in [24].

*Step 2. Reference sample normalization (Normalization I).* For each PSM, the reporter ion intensities are log2 transformed, and the reference sample intensity is subtracted from each sample intensity. Thus, the data is converted, on the PSM-by-PSM basis, to log2-based ratios of intensities with respect to the specified reference (referred to as ratios below). TMT-Integrator supports two approaches for defining the reference for ratio conversion (**Figure 2a**): 1) “Reference sample” approach in which the reference sample is one of the actual samples in the TMT plex. Typically, it would be the common (often referred to as “bridge”) sample used in every TMT plex to assist with normalizing peptide and protein abundances across multiple TMT plexes. In the case of the ccRCC dataset, for example, the reference sample was created by pooling all samples in the study. 2) “Virtual reference” approach, useful when no reference sample is available or specified by the user, in which case the normalization is done with respect to an internally created virtual reference intensity. By default, the virtual reference intensity is calculated for each PSM as the median of all reporter ion intensities in the corresponding MS/MS spectrum. In addition, there is an optional, additional PSM normalization by retention time available, which is not used in any default FragPipe workflows. Specifically, after conversion of intensities to ratios, each channel (except the reference channel when used for normalization) in a PSM table can be divided into ten retention time bins, and the median ratio in the corresponding bin is then subtracted from each individual ratio.

*Step 3. Grouping.* After filtering, reference normalization (conversion to ratios), and optional retention time-based normalization, the selected PSMs are grouped at various levels (gene and protein, but also peptide and modification site-level if PTM-level reports are requested by the user), see **Figure 2b**. At the protein level, PSMs are grouped based on the protein to which they are assigned as unique or razor peptides. At the gene level, PSMs are grouped based on the gene symbol of the corresponding protein to which they were assigned as either unique or razor peptides. Note that, when PTM reports are requested, the gene and protein level tables are based only on the PSMs containing the user specified PTM (e.g., in the case of phosphorylation, the protein level table reports abundance of proteins in the phosphorylated form only). To generate peptide-level and site-level tables (again, for the PTM of interest specified by the user), additional post-processing is applied to generate all non-conflicting PTM configurations (of the specified PTM type) using the strategy similar to that described in [61] (**Figure 1c; Supplementary** Figure 1). In phosphopeptide-enriched datasets, or other PTMs when PTMProphet is used for site localization, confidently localized sites are defined as sites with PTMProphet localization probability of 0.75 or higher. In the table, same peptide sequences but with different PTM site configurations (different site phosphorylation localization configurations or peptides with unlocalized sites), are first indexed as separate entries (these tables are referred to as “multi-site”) Additional, “single-site” tables will be described below. In the peptide-level tables, different site-level configurations are combined into a single peptide-level index, grouping PSMs with all site configurations together if they corresponded to the same peptide sequence. When PTMProphet is not used for computing the site localization probabilities, the reports are generated based on the site assignment from the MSFragger search engine.

*Step 4. Outlier Removal.* In the integrated table, an entry (i.e., a gene/protein/ modified peptide/modified site) may have multiple PSMs identified and grouped to that entry in a particular sample. To select a reasonable abundance ratio to represent the entry, the interquartile range (IQR) algorithm is first applied for outlier removal in each PSM group. We calculate the first quantile (Q1), the third quantile (Q3), and the interquartile range (IQR, i.e., Q3-Q1) of channel ratios. PSMs with ratios outside of the boundaries of Q1-1.5*IQR and Q3+1.5*IQR are excluded from calculating the ratio. The outlier removal step is performed independently at each level, and only for groups containing at least four PSMs.

*Step 5. Abundance Aggregation.* TMT-Integrator provides two abundance aggregation methods, median (default) and weighted median. The median method is a common approach that takes the median ratio of all PSMs (after outlier removal) for the same protein (or peptide, or gene, or multi-site) as the protein (peptide/gene/ multi-site) ratio. The weighted median ratio aggregation method resembles that used in MaxQuant [37], and is implemented as follows. First, the precursor intensity for each PSM in a PSM group is exponentiated, generating a weighting factor for each PSM. The PSMs are then sorted using the ratio to reference in the ascending order. A cumulative weight is incremented by the weighting factor along the sorted PSMs. The ratio where the cumulative weight exceeds 0.5 is taken as the representative of the group, otherwise the ratio with highest weighting factor will be selected. The representative ratios are used as the final values at the corresponding level of grouping (gene, protein, peptide, etc.).

*Step 6. Normalization II.* Two alternative normalization methods, conventional sample-specific median centering (MD) and global normalization (GN), are used to normalize the ratios in the integrated tables, separately at each level of data summarization (gene, protein, peptide, multi-site, single-site). Considering the p (number of entries, e.g. proteins) by n (number of samples) table of ratios (with the samples used as the reference samples excluded), for each entry j in sample i, R_ij_, the median ratio is computed, M_i_ = median(R_ij_, j=1,…,p). The ratios in each sample are then median centered, R^MD^_ij_ = R_ij_ – M_i_. As a result, in the MD ratio tables, the median (log2 scale) ratio for each sample is zero. To generate the alternative, GN-normalized ratio tables, the median absolute deviation (MAD) of the median centered values in each sample are calculated, MAD_i_ = median(abs(R^MD^_ij_), j=1…p), along with the global absolute deviation, MAD_0_=median(MAD_i_, i=1,…,n). All median centered ratios are then additionally scaled to equalize the ratio distributions across all samples, R^GN^_ij_ = (R^MD^_ij_/ MAD_i_) × MAD_0_. Median centering is used as the default approach.

*Step 7. Conversion of ratios back to absolute intensities (abundances)*. As the last step, the normalized ratios (unnormalized, MD, or GN normalized) are converted back to the absolute intensity scale using the estimated intensity of each entry (at each level, gene, protein, peptide, multi-site) in the reference sample. By default (controlled by “use MS1 intensity” option in FragPipe), the absolute intensity of the entry i measured in TMT plex k (k=1,…,q), Ref_ik_, is estimated using the weighted sum of the MS1 intensities of the top 3 most intense peptide ions [62] quantified for that entry in the TMT plex k. The weighting factor, for each PSM, is taken as the proportion of the reference channel TMT intensity to the total summed TMT channel intensity. The overall (based on all q TMT plexes in the dataset) reference intensity for entry i is then estimated as Ref_i_ = median(Ref_ik_, k = 1,…,q). In doing so, the missing intensity values (i.e., no identified and or quantified PSMs for that entry in a particular TMT plex) are imputed with a global minimum intensity value. The final intensity of entry i in sample j (log2 scale) is computed as A_ij_ = R^MD^_ij_ + log_2_(Ref_i_) in the case of MD-normalized ratios (and similarly for the unnormalized or GN-normalized tables). TMT-Integrator also provides an option to perform conversion of ratios to absolute intensities using the total summed reporter ion intensity in MS2 spectra in place of the MS1 precursor intensity (“use MS1 intensity” option unchecked).

In PTM studies, TMT-Integrator also provides single-site PTM reports generated by additional processing of the multi-site reports described above (**Figure 1c; Supplementary** Figure 1), illustrated here using phosphorylation as example. Single-site specific representation of the quantitative PTM data is necessary for some downstream analysis tools, such as PTM-SEA [63]. First, the entries in the multi-site reports that do not contain any localized sites are removed. A new index format with the combination of the protein accession number, modified amino acid, and the localized site position within the protein sequence is created. The entries in the multi-site report are re-clustered using this new single-site index. If a localized site is observed in a singly phosphorylated form (or only in one entry), TMT-Integrator propagates the ratio for that entry to the single-site table. If a localized site is observed on a peptide in singly and doubly phosphorylated forms, TMT-Integrator reports the ratio observed in the monophosphorylated form. If a localized site is observed in double or triple phosphorylated forms, TMT-Integrator reports the median ratios. As for other tables, TMT-Integrator also converts the ratios back to the absolute intensities, using the same logic. For example, localized sites on peptides in singly phosphorylated forms inherit intensities directly from the corresponding entries in the multi-site table. For localized sites without singly phosphorylated forms (i.e., those found only on doubly or triply phosphorylated peptides), the median intensity of those multiply phosphorylated forms is used to estimate the site-level intensity.

When generating these final matrices, an additional filter is applied in TMT-Integrator to remove entries for which the best (highest probability, across the entire dataset) supporting PSM has the probability below a certain threshold (allowing application of an even higher stringency of filtering than the FDR thresholds applied in Philosopher filter command at the earlier stage; controlled by “min best peptide probability” parameter in FragPipe/TMT-Integrator).

## DATA AVAILABILITY

All the data used in this study is publicly available and can be downloaded from the National Cancer Institute Proteomic Data Common (PDC, https://pdc.cancer.gov/pdc/) under study PDC000127 and PDC000128; from ProteomeXchange under accession code: PXD005486, PXD058918 and PXD054559. The parameter and results files generated in this study can be found at https://doi.org/10.5281/zenodo.14983580.

## CODE AVAILABILITY

TMT-Integrator are freely available and can be downloaded at https://github.com/Nesvilab/TMT-Integrator. The R scripts for processing the results and generating the figures are available at https://github.com/Nesvilab/TMT-Integrator-manuscript.

## ACKNOWLEDGEMENT

This work was supported in part by the National Institutes of Health grants R01-GM-094231 and U2CES030164, and the Ministry of Science and Technology of Taiwan (MOST-110-2320-B-008-001-MY2).

