## Supplementary File 1 for "Analysis of isobaric quantitative proteomic data using TMT-Integrator and FragPipe computational platform"

Omics data tables evaluation report


### Omics data tables evaluation report

#### 2024-10-18

### 1 Introduction

In this evaluation, there are a total of **9** data tables. Evaluation metrics from the **OmicsEV** package for these data tables are included in this report, beginning with a summary of the data. The sample distribution by class for each data table is shown in the table below.

| class | FP-RefRatio | FP-RefRatio-MD | FP-RefRatio-Weighted-MD | FP-VirtualRatio-MD | MQ-NoNorm | MQ-NoNorm-MD | MQ-RefRatio-MD | MQ-RefRatio-Weighted-MD | MQ-VirtualRatio-MD |
| --- | --- | --- | --- | --- | --- | --- | --- | --- | --- |
| Normal | 79 | 79 | 79 | 79 | 79 | 79 | 79 | 79 | 79 |
| Tumor | 103 | 103 | 103 | 103 | 103 | 103 | 103 | 103 | 103 |

Detailed information for each sample included in all data tables is shown below.

| sample | class | batch | order |
| --- | --- | --- | --- |
| C3L-01287-N | Normal | 1 | 1 |
| C3L-00561-N | Normal | 1 | 2 |
| C3L-01603-N | Normal | 1 | 3 |
| C3N-00834-N | Normal | 1 | 4 |
| C3N-01214-N | Normal | 1 | 5 |
| C3N-01261-N | Normal | 1 | 6 |
| C3L-00917-N | Normal | 1 | 7 |
| C3L-00607-N | Normal | 1 | 8 |
| C3N-00194-N | Normal | 1 | 9 |
| C3L-00010-N | Normal | 1 | 10 |
| C3L-01861-N | Normal | 1 | 11 |
| C3N-01646-N | Normal | 1 | 12 |
| C3L-01281-N | Normal | 1 | 13 |
| C3N-00495-N | Normal | 1 | 14 |
| C3N-00831-N | Normal | 1 | 15 |
| C3L-00183-N | Normal | 1 | 16 |
| C3N-00168-N | Normal | 1 | 17 |
| C3L-00369-N | Normal | 1 | 18 |
| C3L-00791-N | Normal | 1 | 19 |
| C3L-00097-N | Normal | 1 | 20 |
| C3L-00004-N | Normal | 1 | 21 |
| C3N-00953-N | Normal | 1 | 22 |
| C3N-00150-N | Normal | 1 | 23 |
| C3N-00244-N | Normal | 1 | 24 |
| C3N-01178-N | Normal | 1 | 25 |
| C3L-00583-N | Normal | 1 | 26 |
| C3L-00088-N | Normal | 1 | 27 |
| C3L-00814-N | Normal | 1 | 28 |
| C3L-01885-N | Normal | 1 | 29 |
| C3L-00908-N | Normal | 1 | 30 |
| C3L-00026-N | Normal | 1 | 31 |
| C3L-00447-N | Normal | 1 | 32 |
| C3L-00416-N | Normal | 1 | 33 |
| C3N-00246-N | Normal | 1 | 34 |
| C3N-00148-N | Normal | 1 | 35 |
| C3N-00646-N | Normal | 1 | 36 |
| C3L-00902-N | Normal | 1 | 37 |
| C3L-00096-N | Normal | 1 | 38 |
| C3L-01836-N | Normal | 1 | 39 |
| C3N-00177-N | Normal | 1 | 40 |
| C3N-01200-N | Normal | 1 | 41 |
| C3L-00011-N | Normal | 1 | 42 |
| C3N-01808-N | Normal | 1 | 43 |
| C3N-01176-N | Normal | 1 | 44 |
| C3L-00103-N | Normal | 1 | 45 |
| C3N-01649-N | Normal | 1 | 46 |
| C3N-00149-N | Normal | 1 | 47 |
| C3L-00581-N | Normal | 1 | 48 |
| C3L-01313-N | Normal | 1 | 49 |
| C3N-00573-N | Normal | 1 | 50 |
| C3L-01302-N | Normal | 1 | 51 |
| C3N-01361-N | Normal | 1 | 52 |
| C3N-01651-N | Normal | 1 | 53 |
| C3L-00360-N | Normal | 1 | 54 |
| C3L-00448-N | Normal | 1 | 55 |
| C3L-00907-N | Normal | 1 | 56 |
| C3L-01286-N | Normal | 1 | 57 |
| C3L-01607-N | Normal | 1 | 58 |
| C3N-01179-N | Normal | 1 | 59 |
| C3L-00606-N | Normal | 1 | 60 |
| C3N-01648-N | Normal | 1 | 61 |
| C3N-00242-N | Normal | 1 | 62 |
| C3L-00418-N | Normal | 1 | 63 |
| C3N-00577-N | Normal | 1 | 64 |
| C3L-01882-N | Normal | 1 | 65 |
| C3N-00733-N | Normal | 1 | 66 |
| C3L-00910-N | Normal | 1 | 67 |
| C3L-00079-N | Normal | 1 | 68 |
| C3N-01522-N | Normal | 1 | 69 |
| C3N-01220-N | Normal | 1 | 70 |
| C3N-00852-N | Normal | 1 | 71 |
| C3N-00320-N | Normal | 1 | 72 |
| C3N-00437-N | Normal | 1 | 73 |
| C3N-00491-N | Normal | 1 | 74 |
| C3N-00317-N | Normal | 1 | 75 |
| C3N-00310-N | Normal | 1 | 76 |
| C3N-00390-N | Normal | 1 | 77 |
| C3N-00312-N | Normal | 1 | 78 |
| C3N-00494-N | Normal | 1 | 79 |
| C3N-01648-T | Tumor | 1 | 80 |
| C3N-01214-T | Tumor | 1 | 81 |
| C3N-00646-T | Tumor | 1 | 82 |
| C3L-00766-T | Tumor | 1 | 83 |
| C3L-00360-T | Tumor | 1 | 84 |
| C3L-00790-T | Tumor | 1 | 85 |
| C3N-00154-T | Tumor | 1 | 86 |
| C3L-00765-T | Tumor | 1 | 87 |
| C3N-00150-T | Tumor | 1 | 88 |
| C3L-01283-T | Tumor | 1 | 89 |
| C3N-00315-T | Tumor | 1 | 90 |
| C3N-00177-T | Tumor | 1 | 91 |
| C3L-00561-T | Tumor | 1 | 92 |
| C3L-00800-T | Tumor | 1 | 93 |
| C3L-01882-T | Tumor | 1 | 94 |
| C3N-00491-T | Tumor | 1 | 95 |
| C3N-00577-T | Tumor | 1 | 96 |
| C3L-00079-T | Tumor | 1 | 97 |
| C3N-01220-T | Tumor | 1 | 98 |
| C3L-01607-T | Tumor | 1 | 99 |
| C3N-00953-T | Tumor | 1 | 100 |
| C3N-00380-T | Tumor | 1 | 101 |
| C3N-00317-T | Tumor | 1 | 102 |
| C3N-00305-T | Tumor | 1 | 103 |
| C3N-00437-T | Tumor | 1 | 104 |
| C3N-01651-T | Tumor | 1 | 105 |
| C3L-00908-T | Tumor | 1 | 106 |
| C3L-00416-T | Tumor | 1 | 107 |
| C3N-00733-T | Tumor | 1 | 108 |
| C3L-00610-T | Tumor | 1 | 109 |
| C3N-01213-T | Tumor | 1 | 110 |
| C3N-01200-T | Tumor | 1 | 111 |
| C3N-01361-T | Tumor | 1 | 112 |
| C3L-00183-T | Tumor | 1 | 113 |
| C3L-01861-T | Tumor | 1 | 114 |
| C3L-00917-T | Tumor | 1 | 115 |
| C3L-00817-T | Tumor | 1 | 116 |
| C3L-00581-T | Tumor | 1 | 117 |
| C3L-00004-T | Tumor | 1 | 118 |
| C3N-01524-T | Tumor | 1 | 119 |
| C3L-01302-T | Tumor | 1 | 120 |
| C3L-01836-T | Tumor | 1 | 121 |
| C3N-00314-T | Tumor | 1 | 122 |
| C3L-01287-T | Tumor | 1 | 123 |
| C3N-00149-T | Tumor | 1 | 124 |
| C3N-00148-T | Tumor | 1 | 125 |
| C3N-00390-T | Tumor | 1 | 126 |
| C3L-01885-T | Tumor | 1 | 127 |
| C3N-00495-T | Tumor | 1 | 128 |
| C3N-01646-T | Tumor | 1 | 129 |
| C3L-00813-T | Tumor | 1 | 130 |
| C3L-00812-T | Tumor | 1 | 131 |
| C3N-00494-T | Tumor | 1 | 132 |
| C3N-00573-T | Tumor | 1 | 133 |
| C3L-01560-T | Tumor | 1 | 134 |
| C3N-01649-T | Tumor | 1 | 135 |
| C3L-00792-T | Tumor | 1 | 136 |
| C3N-01179-T | Tumor | 1 | 137 |
| C3N-00168-T | Tumor | 1 | 138 |
| C3L-00607-T | Tumor | 1 | 139 |
| C3L-00907-T | Tumor | 1 | 140 |
| C3N-01178-T | Tumor | 1 | 141 |
| C3L-00902-T | Tumor | 1 | 142 |
| C3N-00831-T | Tumor | 1 | 143 |
| C3L-00096-T | Tumor | 1 | 144 |
| C3N-00194-T | Tumor | 1 | 145 |
| C3L-00010-T | Tumor | 1 | 146 |
| C3N-00852-T | Tumor | 1 | 147 |
| C3N-01522-T | Tumor | 1 | 148 |
| C3L-01288-T | Tumor | 1 | 149 |
| C3L-00799-T | Tumor | 1 | 150 |
| C3L-00088-T | Tumor | 1 | 151 |
| C3L-00910-T | Tumor | 1 | 152 |
| C3N-01261-T | Tumor | 1 | 153 |
| C3L-00606-T | Tumor | 1 | 154 |
| C3L-01603-T | Tumor | 1 | 155 |
| C3L-01553-T | Tumor | 1 | 156 |
| C3L-01281-T | Tumor | 1 | 157 |
| C3N-00320-T | Tumor | 1 | 158 |
| C3L-01557-T | Tumor | 1 | 159 |
| C3L-01352-T | Tumor | 1 | 160 |
| C3N-01176-T | Tumor | 1 | 161 |
| C3N-00834-T | Tumor | 1 | 162 |
| C3L-00418-T | Tumor | 1 | 163 |
| C3L-00796-T | Tumor | 1 | 164 |
| C3L-00103-T | Tumor | 1 | 165 |
| C3L-00097-T | Tumor | 1 | 166 |
| C3N-00312-T | Tumor | 1 | 167 |
| C3L-00583-T | Tumor | 1 | 168 |
| C3L-00814-T | Tumor | 1 | 169 |
| C3L-01313-T | Tumor | 1 | 170 |
| C3L-00026-T | Tumor | 1 | 171 |
| C3L-00447-T | Tumor | 1 | 172 |
| C3L-00448-T | Tumor | 1 | 173 |
| C3N-01808-T | Tumor | 1 | 174 |
| C3L-00011-T | Tumor | 1 | 175 |
| C3N-00310-T | Tumor | 1 | 176 |
| C3N-00244-T | Tumor | 1 | 177 |
| C3L-00791-T | Tumor | 1 | 178 |
| C3L-01286-T | Tumor | 1 | 179 |
| C3N-00246-T | Tumor | 1 | 180 |
| C3N-00242-T | Tumor | 1 | 181 |
| C3L-00369-T | Tumor | 1 | 182 |

### 2 Overview

The table below provides an overview about all the quantitative metrics generated in the evaluation. For each metric, the value of the best data table is highlighted in bold and red. The details for each metric can be found in the corresponding sections below.

| metric | FP-RefRatio | FP-RefRatio-MD | FP-RefRatio-Weighted-MD | FP-VirtualRatio-MD | MQ-NoNorm | MQ-NoNorm-MD | MQ-RefRatio-MD | MQ-RefRatio-Weighted-MD | MQ-VirtualRatio-MD |
| --- | --- | --- | --- | --- | --- | --- | --- | --- | --- |
| #identified features | 12210 (0.5989) | 12210 (0.5989) | 12210 (0.5989) | 12210 (0.5989) | 10815 (0.5305) | 10815 (0.5305) | 10815 (0.5305) | 10815 (0.5305) | 10815 (0.5305) |
| #quantifiable features | 9521 (0.4670) | 9521 (0.4670) | 9521 (0.4670) | 9521 (0.4670) | 8463 (0.4151) | 8463 (0.4151) | 8462 (0.4151) | 8462 (0.4151) | 8463 (0.4151) |
| non\_missing\_value\_ratio | 0.9521 | 0.9521 | 0.9521 | 0.9521 | 0.9397 | 0.9397 | 0.9397 | 0.9397 | 0.9397 |
| data\_dist\_similarity | 0.9526 | 0.9823 | 0.9820 | 0.9812 | 0.9099 | 0.9959 | 0.9752 | 0.9840 | 0.9763 |
| complex\_auc | 0.8167 | 0.8601 | 0.8533 | 0.8715 | 0.6487 | 0.6227 | 0.8435 | 0.7156 | 0.8617 |
| func\_auc | 0.8353 | 0.8276 | 0.8202 | 0.8401 | 0.6618 | 0.7050 | 0.7085 | 0.7385 | 0.7467 |
| gene\_wise\_cor | 0.2792 | 0.4050 | 0.3939 | 0.3720 | 0.1875 | 0.2127 | 0.4071 | 0.2625 | 0.3667 |
| sample\_wise\_cor | 0.4641 | 0.4641 | 0.4664 | 0.4594 | 0.4148 | 0.4148 | 0.1856 | 0.3047 | 0.1693 |

The radar plot below summarizes results from the overview table above. To generate the radar plot, each metric is scaled from 0 to 1 such that higher values indicate better data quality if necessary. Scaled values are in parentheses in the table.

### 3 Data depth

#### 3.1 Study-wise

The table below shows the number of identified and quantified proteins or genes for each data table. Identified proteins or genes are those with a measurement in any sample in a data table whereas quantified proteins or genes are those that remain after filtering out those with missing values in more than 50% of the samples in a data table. The values in parentheses are the percentage of proteins or genes identified or quantified based on the total number of proteins or genes (20386) in the study species.

| data table | #identified features | #quantifiable features |
| --- | --- | --- |
| FP-RefRatio-MD | 12210 (59.89%) | 9521 (46.70%) |
| FP-RefRatio-Weighted-MD | 12210 (59.89%) | 9521 (46.70%) |
| FP-RefRatio | 12210 (59.89%) | 9521 (46.70%) |
| FP-VirtualRatio-MD | 12210 (59.89%) | 9521 (46.70%) |
| MQ-NoNorm-MD | 10815 (53.05%) | 8463 (41.51%) |
| MQ-NoNorm | 10815 (53.05%) | 8463 (41.51%) |
| MQ-RefRatio-MD | 10815 (53.05%) | 8462 (41.51%) |
| MQ-RefRatio-Weighted-MD | 10815 (53.05%) | 8462 (41.51%) |
| MQ-VirtualRatio-MD | 10815 (53.05%) | 8463 (41.51%) |

The upset chart below shows overlap between proteins or genes identified in each data table. Numbers of proteins or genes commonly identified in different combinations of data tables are indicated in the top bar chart, and the specific combinations of data tables containing those proteins or genes are indicated with solid points below the bar chart. Total identifications for each data table are indicated on the right as ‘Set size’.

#### 3.2 Sample-wise

The figures below show the number of proteins or genes identified/quantified (non-missing values) in each sample. Samples from different batches are coded with different shapes, and samples from different classes are coded with different colors. A separate figure is shown for each data table.

FP-RefRatio-MDFP-RefRatio-Weighted-MDFP-RefRatioFP-VirtualRatio-MDMQ-NoNorm-MDMQ-NoNormMQ-RefRatio-MDMQ-RefRatio-Weighted-MDMQ-VirtualRatio-MD

#### 3.3 Missing value distribution

The missing value distribution provides an overview of the completeness of the data. The table below shows the percent of missing values for all samples in each data table.

| data table | non\_missing\_value\_ratio |
| --- | --- |
| FP-RefRatio-MD | 0.9521 |
| FP-RefRatio-Weighted-MD | 0.9521 |
| FP-RefRatio | 0.9521 |
| FP-VirtualRatio-MD | 0.9521 |
| MQ-NoNorm-MD | 0.9397 |
| MQ-NoNorm | 0.9397 |
| MQ-RefRatio-MD | 0.9397 |
| MQ-RefRatio-Weighted-MD | 0.9397 |
| MQ-VirtualRatio-MD | 0.9397 |

The following barplots show missing value distributions for each data table as number (Y axis)/percentage (number above bar) of proteins or genes with missing values in each bin. Genes are binned by proportion of samples with missing values from 0.1 to 1 in increments of 0.1, where 0.1 indicates missing values in no more than 10% of the samples, and 1 indicates missing values in all samples.

FP-RefRatio-MDFP-RefRatio-Weighted-MDFP-RefRatioFP-VirtualRatio-MDMQ-NoNorm-MDMQ-NoNormMQ-RefRatio-MDMQ-RefRatio-Weighted-MDMQ-VirtualRatio-MD

### 4 Data normalization

#### 4.1 Boxplot

Normalized data is expected to be centered around a similar value and show similar distributions in all samples. The boxplots below show the protein or gene expression measurement distribution across samples in each data table, allowing for qualitative assessment of the normalized data. Samples in input order are indicated on the X axis. The Y axis shows log2 transformed protein or gene values. Samples from different classes are coded with different colors.

FP-RefRatio-MDFP-RefRatio-Weighted-MDFP-RefRatioFP-VirtualRatio-MDMQ-NoNorm-MDMQ-NoNormMQ-RefRatio-MDMQ-RefRatio-Weighted-MDMQ-VirtualRatio-MD

To quantify the normalization effect, we tested for how well the data in the feature set can distinguish between each pair of samples. If the distribution is similar for the two samples in a given pair, the overall feature abundance (levels for all features in one sample vs the other) should not be sufficient to predict which sample is which. Therefore, for each pair of samples, an AUROC test was performed to quantify the ability of feature abundance to distinguish the two samples, and then a **data\_dist\_similarity** score was generated: 1-2\*abs(AUROC-0.5). This score ranges from 0 to 1, and the higher the score is the better the normalized data quality is (no systematic difference between the two samples). The final metric for each data table is the median of scores from all sample pairs. The column ‘n’ shows the total number of sample pairs in the analysis.

| data table | data\_dist\_similarity | n |
| --- | --- | --- |
| FP-RefRatio-MD | 0.9823 | 16471 |
| FP-RefRatio-Weighted-MD | 0.9820 | 16471 |
| FP-RefRatio | 0.9526 | 16471 |
| FP-VirtualRatio-MD | 0.9812 | 16471 |
| MQ-NoNorm-MD | 0.9959 | 16471 |
| MQ-NoNorm | 0.9099 | 16471 |
| MQ-RefRatio-MD | 0.9752 | 16471 |
| MQ-RefRatio-Weighted-MD | 0.9840 | 16471 |
| MQ-VirtualRatio-MD | 0.9763 | 16471 |

#### 4.2 Density plot

The density plots below show the expression distributions for all samples (separate line) in each data table. The Y axis shows the density over the range of log2 transformed protein or gene expression values (X axis).

### 5 Batch effect

#### 5.1 Correlation heatmap

Another way to qualitatively assess batch effect is to visualize the correlations for measurements between samples from the same batch to those in samples from different batches using heatmaps. The following figures show Spearman correlation heatmaps for all pairs of samples (all samples included in both rows and columns) for each data table. The color indicates the correlation between samples. The samples are ordered by batches. Concentration of high correlation values (red color) for pairs of samples from the same batch block compared to other batches indicates the presence of batch effect.

FP-RefRatio-MDFP-RefRatio-Weighted-MDFP-RefRatioFP-VirtualRatio-MDMQ-NoNorm-MDMQ-NoNormMQ-RefRatio-MDMQ-RefRatio-Weighted-MDMQ-VirtualRatio-MD

### 6 Biological signal

#### 6.1 Correlation among protein complex members

Members of the same protein complex often show greater correlation in gene and protein expression (IntraComplex correlation) than genes or proteins that are in different complexes (InterComplex correlation). Thus, one way to evaluate the quality of the biological signal present in a data table is to compare IntraComplex correlation to InterComplex correlation. Furthermore, because of the need to preserve stoichiometry between protein complex members, the difference between IntraComplex correlation and InterComplex correlation is often greater at the protein level than at the RNA data. If both RNA and protein data tables are available, observing that this difference is more pronounced in the protein data table than the RNA data table serves as an indicator for the quality of the protein data. We use the protein complexes from the CORUM database in this analysis.

The boxplots below show the distributions and ranges for pairwise correlations between genes or proteins from the same complex and for genes and proteins from different complexes for each data table.

The table below shows a summary of the evaluation. ‘diff’ is Cor(intra) - Cor(inter). ‘complex\_auc’ is the AUROC value based on correlation of protein pairs from different groups.

| data table | InterComplex | IntraComplex | diff | complex\_auc |
| --- | --- | --- | --- | --- |
| FP-RefRatio | 0.6583 | 0.8656 | 0.2073 | 0.8167 |
| FP-RefRatio-MD | 0.0150 | 0.4244 | 0.4094 | 0.8601 |
| FP-RefRatio-Weighted-MD | 0.0188 | 0.3935 | 0.3746 | 0.8533 |
| FP-VirtualRatio-MD | -0.0027 | 0.4916 | 0.4943 | 0.8715 |
| MQ-NoNorm | 0.3326 | 0.4477 | 0.1151 | 0.6487 |
| MQ-NoNorm-MD | 0.0210 | 0.1100 | 0.0890 | 0.6227 |
| MQ-RefRatio-MD | 0.0077 | 0.3613 | 0.3535 | 0.8435 |
| MQ-RefRatio-Weighted-MD | 0.0058 | 0.1372 | 0.1314 | 0.7156 |
| MQ-VirtualRatio-MD | -0.0048 | 0.4401 | 0.4449 | 0.8617 |
| RNA | 0.0163 | 0.1472 | 0.1310 | 0.6570 |

#### 6.2 Gene function prediction

Previous studies have shown that expression correlation is often higher for functionally related genes or proteins than for unrelated genes or proteins and that this correlation is greater when considering protein data than when considering RNA data (Wang, Jing, et al. Molecular & Cellular Proteomics 16.1 (2017): 121-134.). Therefore, we can also evaluate the biological signal present in a data table by evaluating functional category predictions made using a co-expression network generated from each data table.

In this evaluation, each data table was used to build a co-expression network. For a selected network and a selected functional category (such as a selected category from GO or KEGG), proteins/genes annotated to the category and also included in the network were defined as a positive protein/gene set, and other proteins/genes in the network constituted the negative protein/gene set for the category. For a selected functional category, a subset of the proteins/genes were used as seed proteins/genes for random walk through the network to calculate scores for other proteins/genes. A higher score for a protein/gene represents a closer relationship between the protein/gene and the seed proteins/genes. The table below shows AUROCs of the prediction performance using this score for each selected functional category.

|  | FP-RefRatio | FP-RefRatio-MD | FP-RefRatio-Weighted-MD | FP-VirtualRatio-MD | MQ-NoNorm | MQ-NoNorm-MD | MQ-RefRatio-MD | MQ-RefRatio-Weighted-MD | MQ-VirtualRatio-MD | RNA |
| --- | --- | --- | --- | --- | --- | --- | --- | --- | --- | --- |
| Acute myeloid leukemia | 0.83 | 0.735 | 0.712 | 0.789 | 0.655 | 0.611 | 0.531 | 0.654 | 0.679 | 0.628 |
| Adherens junction | 0.82 | 0.759 | 0.755 | 0.756 | 0.604 | 0.648 | 0.774 | 0.511 | 0.747 | 0.595 |
| Adipocytokine signaling pathway | 0.714 | 0.61 | 0.651 | 0.706 | 0.578 | 0.661 | 0.567 | 0.669 | 0.615 | 0.545 |
| African trypanosomiasis | 0.688 | 0.633 | 0.705 | 0.652 | 0.693 | 0.613 | 0.753 | 0.758 | 0.777 | 0.751 |
| Allograft rejection | 0.929 | 0.988 | 0.941 | 0.987 | 0.846 | 0.783 | 0.962 | 0.975 | 0.971 | 0.964 |
| Alzheimers disease | 0.819 | 0.828 | 0.837 | 0.868 | 0.563 | 0.641 | 0.599 | 0.72 | 0.562 | 0.776 |
| Amino sugar and nucleotide sugar metabolism | 0.711 | 0.752 | 0.784 | 0.727 | 0.569 | 0.58 | 0.775 | 0.575 | 0.776 | 0.68 |
| Amoebiasis | 0.772 | 0.781 | 0.795 | 0.783 | 0.689 | 0.665 | 0.76 | 0.711 | 0.746 | 0.742 |
| Amyotrophic lateral sclerosis (ALS) | 0.636 | 0.603 | 0.563 | 0.531 | 0.559 | 0.63 | 0.57 | 0.629 | 0.605 | 0.605 |
| Antigen processing and presentation | 0.96 | 0.97 | 0.933 | 0.959 | 0.663 | 0.732 | 0.958 | 0.811 | 0.919 | 0.895 |
| Apoptosis | 0.703 | 0.717 | 0.667 | 0.746 | 0.567 | 0.576 | 0.626 | 0.578 | 0.707 | 0.593 |
| Arachidonic acid metabolism | 0.68 | 0.579 | 0.582 | 0.61 | 0.631 | 0.595 | 0.558 | 0.748 | 0.603 | 0.61 |
| Arginine and proline metabolism | 0.868 | 0.857 | 0.838 | 0.787 | 0.633 | 0.729 | 0.672 | 0.785 | 0.726 | 0.52 |
| Arrhythmogenic right ventricular cardiomyopathy (ARVC) | 0.715 | 0.744 | 0.765 | 0.841 | 0.637 | 0.663 | 0.659 | 0.616 | 0.768 | 0.664 |
| Autoimmune thyroid disease | 0.91 | 0.988 | 0.942 | 0.984 | 0.76 | 0.805 | 0.981 | 0.977 | 0.973 | 0.964 |
| Axon guidance | 0.701 | 0.75 | 0.649 | 0.729 | 0.572 | 0.604 | 0.617 | 0.594 | 0.596 | 0.599 |
| B cell receptor signaling pathway | 0.74 | 0.773 | 0.764 | 0.78 | 0.632 | 0.697 | 0.665 | 0.683 | 0.715 | 0.58 |
| Bacterial invasion of epithelial cells | 0.776 | 0.789 | 0.746 | 0.786 | 0.582 | 0.58 | 0.701 | 0.631 | 0.756 | 0.623 |
| Bile secretion | 0.526 | 0.855 | 0.832 | 0.767 | 0.662 | 0.605 | 0.732 | 0.689 | 0.7 | 0.561 |
| Bladder cancer | 0.757 | 0.638 | 0.666 | 0.684 | 0.561 | 0.59 | 0.587 | 0.609 | 0.654 | 0.552 |
| Calcium signaling pathway | 0.785 | 0.689 | 0.63 | 0.699 | 0.642 | 0.625 | 0.706 | 0.593 | 0.772 | 0.567 |
| Carbohydrate digestion and absorption | 0.668 | 0.706 | 0.732 | 0.705 | 0.648 | 0.636 | 0.857 | 0.718 | 0.577 | 0.65 |
| Cell adhesion molecules (CAMs) | 0.835 | 0.808 | 0.785 | 0.894 | 0.596 | 0.628 | 0.807 | 0.811 | 0.796 | 0.741 |
| Cell cycle | 0.809 | 0.795 | 0.787 | 0.855 | 0.667 | 0.701 | 0.808 | 0.718 | 0.817 | 0.772 |
| Chagas disease (American trypanosomiasis) | 0.746 | 0.748 | 0.799 | 0.756 | 0.584 | 0.567 | 0.721 | 0.527 | 0.724 | 0.551 |
| Chemokine signaling pathway | 0.779 | 0.714 | 0.734 | 0.735 | 0.62 | 0.679 | 0.674 | 0.584 | 0.682 | 0.654 |
| Chronic myeloid leukemia | 0.727 | 0.738 | 0.689 | 0.748 | 0.582 | 0.578 | 0.69 | 0.648 | 0.676 | 0.519 |
| Colorectal cancer | 0.767 | 0.649 | 0.602 | 0.732 | 0.62 | 0.617 | 0.738 | 0.585 | 0.698 | 0.583 |
| Complement and coagulation cascades | 0.95 | 0.95 | 0.971 | 0.92 | 0.94 | 0.937 | 0.881 | 0.984 | 0.951 | 0.774 |
| Cysteine and methionine metabolism | 0.799 | 0.762 | 0.788 | 0.732 | 0.729 | 0.659 | 0.761 | 0.722 | 0.711 | 0.646 |
| Cytokine-cytokine receptor interaction | 0.831 | 0.821 | 0.766 | 0.823 | 0.615 | 0.659 | 0.684 | 0.811 | 0.688 | 0.609 |
| Cytosolic DNA-sensing pathway | 0.638 | 0.646 | 0.679 | 0.709 | 0.59 | 0.58 | 0.712 | 0.693 | 0.534 | 0.525 |
| Dilated cardiomyopathy | 0.745 | 0.777 | 0.803 | 0.803 | 0.602 | 0.65 | 0.706 | 0.59 | 0.767 | 0.7 |
| Drug metabolism - cytochrome P450 | 0.871 | 0.797 | 0.754 | 0.675 | 0.718 | 0.605 | 0.901 | 0.74 | 0.805 | 0.642 |
| ECM-receptor interaction | 0.874 | 0.804 | 0.813 | 0.861 | 0.751 | 0.796 | 0.787 | 0.809 | 0.803 | 0.772 |
| Endocytosis | 0.843 | 0.782 | 0.749 | 0.789 | 0.588 | 0.572 | 0.635 | 0.584 | 0.628 | 0.614 |
| Endometrial cancer | 0.754 | 0.687 | 0.724 | 0.805 | 0.661 | 0.571 | 0.618 | 0.575 | 0.611 | 0.646 |
| Epithelial cell signaling in Helicobacter pylori infection | 0.786 | 0.693 | 0.799 | 0.761 | 0.626 | 0.653 | 0.636 | 0.709 | 0.662 | 0.607 |
| ErbB signaling pathway | 0.755 | 0.708 | 0.633 | 0.683 | 0.597 | 0.534 | 0.621 | 0.634 | 0.605 | 0.604 |
| Fc epsilon RI signaling pathway | 0.836 | 0.727 | 0.783 | 0.808 | 0.59 | 0.624 | 0.787 | 0.674 | 0.739 | 0.64 |
| Fc gamma R-mediated phagocytosis | 0.795 | 0.783 | 0.715 | 0.789 | 0.62 | 0.706 | 0.769 | 0.672 | 0.826 | 0.623 |
| Focal adhesion | 0.774 | 0.787 | 0.789 | 0.782 | 0.69 | 0.738 | 0.754 | 0.696 | 0.776 | 0.746 |
| Fructose and mannose metabolism | 0.795 | 0.797 | 0.777 | 0.758 | 0.649 | 0.726 | 0.801 | 0.724 | 0.797 | 0.658 |
| Galactose metabolism | 0.68 | 0.679 | 0.704 | 0.698 | 0.652 | 0.577 | 0.706 | 0.667 | 0.648 | 0.658 |
| Gap junction | 0.756 | 0.724 | 0.76 | 0.676 | 0.66 | 0.623 | 0.626 | 0.638 | 0.59 | 0.509 |
| Gastric acid secretion | 0.71 | 0.63 | 0.613 | 0.634 | 0.647 | 0.636 | 0.624 | 0.628 | 0.712 | 0.545 |
| Glioma | 0.715 | 0.647 | 0.638 | 0.703 | 0.577 | 0.591 | 0.67 | 0.619 | 0.628 | 0.662 |
| Glutathione metabolism | 0.65 | 0.662 | 0.588 | 0.61 | 0.651 | 0.626 | 0.74 | 0.658 | 0.689 | 0.604 |
| Glycerolipid metabolism | 0.685 | 0.742 | 0.666 | 0.75 | 0.658 | 0.584 | 0.723 | 0.837 | 0.72 | 0.64 |
| Glycerophospholipid metabolism | 0.576 | 0.773 | 0.658 | 0.664 | 0.607 | 0.607 | 0.554 | 0.639 | 0.705 | 0.602 |
| Glycolysis / Gluconeogenesis | 0.853 | 0.819 | 0.784 | 0.863 | 0.669 | 0.722 | 0.837 | 0.769 | 0.817 | 0.712 |
| GnRH signaling pathway | 0.611 | 0.74 | 0.602 | 0.656 | 0.589 | 0.598 | 0.623 | 0.598 | 0.7 | 0.558 |
| Graft-versus-host disease | 0.937 | 0.953 | 0.978 | 0.998 | 0.61 | 0.741 | 0.979 | 0.962 | 0.954 | 0.955 |
| Hematopoietic cell lineage | 0.678 | 0.719 | 0.744 | 0.693 | 0.577 | 0.563 | 0.679 | 0.707 | 0.663 | 0.712 |
| Hepatitis C | 0.792 | 0.784 | 0.755 | 0.855 | 0.604 | 0.597 | 0.762 | 0.628 | 0.736 | 0.535 |
| Huntingtons disease | 0.856 | 0.881 | 0.853 | 0.87 | 0.591 | 0.641 | 0.559 | 0.658 | 0.545 | 0.789 |
| Hypertrophic cardiomyopathy (HCM) | 0.733 | 0.701 | 0.745 | 0.772 | 0.662 | 0.627 | 0.712 | 0.607 | 0.734 | 0.613 |
| Inositol phosphate metabolism | 0.689 | 0.588 | 0.68 | 0.618 | 0.583 | 0.661 | 0.625 | 0.685 | 0.717 | 0.636 |
| Insulin signaling pathway | 0.768 | 0.704 | 0.697 | 0.754 | 0.579 | 0.63 | 0.653 | 0.636 | 0.748 | 0.498 |
| Jak-STAT signaling pathway | 0.732 | 0.773 | 0.775 | 0.859 | 0.601 | 0.663 | 0.588 | 0.734 | 0.608 | 0.582 |
| Leishmaniasis | 0.81 | 0.84 | 0.926 | 0.857 | 0.604 | 0.705 | 0.759 | 0.815 | 0.789 | 0.812 |
| Leukocyte transendothelial migration | 0.808 | 0.753 | 0.745 | 0.767 | 0.587 | 0.666 | 0.775 | 0.747 | 0.809 | 0.659 |
| Long-term depression | 0.777 | 0.767 | 0.725 | 0.7 | 0.609 | 0.698 | 0.616 | 0.58 | 0.612 | 0.562 |
| Long-term potentiation | 0.67 | 0.627 | 0.757 | 0.666 | 0.643 | 0.728 | 0.533 | 0.592 | 0.749 | 0.624 |
| Lysosome | 0.909 | 0.904 | 0.892 | 0.871 | 0.643 | 0.715 | 0.717 | 0.702 | 0.82 | 0.758 |
| Malaria | 0.747 | 0.723 | 0.832 | 0.745 | 0.74 | 0.637 | 0.692 | 0.784 | 0.835 | 0.675 |
| MAPK signaling pathway | 0.722 | 0.68 | 0.687 | 0.622 | 0.626 | 0.657 | 0.557 | 0.528 | 0.629 | 0.606 |
| Melanogenesis | 0.672 | 0.739 | 0.639 | 0.676 | 0.62 | 0.57 | 0.733 | 0.62 | 0.711 | 0.648 |
| Melanoma | 0.768 | 0.732 | 0.776 | 0.746 | 0.601 | 0.626 | 0.669 | 0.595 | 0.62 | 0.585 |
| Metabolic pathways | 0.769 | 0.785 | 0.771 | 0.765 | 0.636 | 0.66 | 0.702 | 0.706 | 0.685 | 0.638 |
| Metabolism of xenobiotics by cytochrome P450 | 0.747 | 0.766 | 0.795 | 0.722 | 0.665 | 0.593 | 0.819 | 0.609 | 0.58 | 0.565 |
| mRNA surveillance pathway | 0.842 | 0.853 | 0.85 | 0.785 | 0.789 | 0.678 | 0.612 | 0.666 | 0.656 | 0.705 |
| mTOR signaling pathway | 0.701 | 0.625 | 0.737 | 0.725 | 0.635 | 0.53 | 0.567 | 0.562 | 0.65 | 0.578 |
| N-Glycan biosynthesis | 0.986 | 0.921 | 0.813 | 0.89 | 0.822 | 0.866 | 0.924 | 0.894 | 0.888 | 0.616 |
| Natural killer cell mediated cytotoxicity | 0.762 | 0.729 | 0.781 | 0.776 | 0.607 | 0.665 | 0.701 | 0.771 | 0.752 | 0.627 |
| Neurotrophin signaling pathway | 0.751 | 0.645 | 0.713 | 0.725 | 0.54 | 0.628 | 0.597 | 0.547 | 0.708 | 0.666 |
| NOD-like receptor signaling pathway | 0.697 | 0.749 | 0.778 | 0.778 | 0.6 | 0.551 | 0.601 | 0.66 | 0.735 | 0.596 |
| Non-small cell lung cancer | 0.771 | 0.69 | 0.629 | 0.758 | 0.583 | 0.559 | 0.699 | 0.635 | 0.705 | 0.613 |
| Nucleotide excision repair | 0.874 | 0.942 | 0.945 | 0.813 | 0.581 | 0.727 | 0.857 | 0.635 | 0.823 | 0.739 |
| Oocyte meiosis | 0.804 | 0.784 | 0.784 | 0.8 | 0.639 | 0.706 | 0.631 | 0.586 | 0.681 | 0.622 |
| Osteoclast differentiation | 0.781 | 0.801 | 0.806 | 0.84 | 0.589 | 0.671 | 0.71 | 0.696 | 0.69 | 0.59 |
| p53 signaling pathway | 0.736 | 0.66 | 0.647 | 0.681 | 0.693 | 0.544 | 0.641 | 0.728 | 0.674 | 0.707 |
| Pancreatic cancer | 0.784 | 0.694 | 0.691 | 0.775 | 0.553 | 0.592 | 0.582 | 0.588 | 0.649 | 0.595 |
| Pancreatic secretion | 0.583 | 0.673 | 0.604 | 0.678 | 0.599 | 0.62 | 0.611 | 0.642 | 0.623 | 0.567 |
| Parkinsons disease | 0.887 | 0.883 | 0.859 | 0.902 | 0.621 | 0.743 | 0.614 | 0.738 | 0.649 | 0.827 |
| Pathogenic Escherichia coli infection | 0.863 | 0.691 | 0.739 | 0.779 | 0.632 | 0.595 | 0.708 | 0.59 | 0.665 | 0.606 |
| Pathways in cancer | 0.712 | 0.647 | 0.615 | 0.636 | 0.562 | 0.57 | 0.532 | 0.538 | 0.605 | 0.568 |
| Pentose phosphate pathway | 0.692 | 0.715 | 0.8 | 0.872 | 0.737 | 0.682 | 0.858 | 0.721 | 0.754 | 0.724 |
| Peroxisome | 0.825 | 0.844 | 0.808 | 0.788 | 0.716 | 0.703 | 0.831 | 0.619 | 0.726 | 0.747 |
| Phagosome | 0.833 | 0.835 | 0.851 | 0.858 | 0.622 | 0.692 | 0.722 | 0.735 | 0.79 | 0.722 |
| Phosphatidylinositol signaling system | 0.659 | 0.578 | 0.629 | 0.581 | 0.592 | 0.64 | 0.65 | 0.596 | 0.601 | 0.706 |
| PPAR signaling pathway | 0.729 | 0.668 | 0.617 | 0.745 | 0.641 | 0.668 | 0.673 | 0.598 | 0.575 | 0.644 |
| Prion diseases | 0.851 | 0.828 | 0.747 | 0.767 | 0.748 | 0.698 | 0.706 | 0.745 | 0.828 | 0.605 |
| Progesterone-mediated oocyte maturation | 0.724 | 0.8 | 0.724 | 0.799 | 0.662 | 0.71 | 0.686 | 0.614 | 0.658 | 0.634 |
| Prostate cancer | 0.742 | 0.6 | 0.691 | 0.677 | 0.596 | 0.572 | 0.563 | 0.549 | 0.7 | 0.607 |
| Proteasome | 0.97 | 0.993 | 0.991 | 0.964 | 0.705 | 0.769 | 0.95 | 0.842 | 0.912 | 0.882 |
| Protein digestion and absorption | 0.848 | 0.818 | 0.828 | 0.883 | 0.753 | 0.878 | 0.77 | 0.837 | 0.845 | 0.689 |
| Protein export | 0.894 | 0.97 | 0.968 | 0.86 | 0.678 | 0.767 | 0.95 | 0.809 | 0.914 | 0.819 |
| Protein processing in endoplasmic reticulum | 0.808 | 0.82 | 0.847 | 0.818 | 0.632 | 0.725 | 0.768 | 0.711 | 0.76 | 0.756 |
| Purine metabolism | 0.623 | 0.753 | 0.689 | 0.662 | 0.56 | 0.567 | 0.718 | 0.524 | 0.647 | 0.575 |
| Pyrimidine metabolism | 0.587 | 0.607 | 0.587 | 0.666 | 0.576 | 0.559 | 0.695 | 0.563 | 0.62 | 0.584 |
| Regulation of actin cytoskeleton | 0.743 | 0.78 | 0.728 | 0.754 | 0.562 | 0.609 | 0.673 | 0.592 | 0.744 | 0.681 |
| Renal cell carcinoma | 0.593 | 0.642 | 0.628 | 0.704 | 0.539 | 0.557 | 0.605 | 0.65 | 0.637 | 0.607 |
| Rheumatoid arthritis | 0.915 | 0.954 | 0.955 | 0.934 | 0.617 | 0.758 | 0.922 | 0.878 | 0.917 | 0.717 |
| Ribosome | 0.969 | 0.96 | 0.965 | 0.98 | 0.698 | 0.829 | 0.935 | 0.856 | 0.956 | 0.931 |
| Ribosome biogenesis in eukaryotes | 0.899 | 0.899 | 0.915 | 0.861 | 0.748 | 0.816 | 0.851 | 0.775 | 0.829 | 0.751 |
| RIG-I-like receptor signaling pathway | 0.729 | 0.714 | 0.737 | 0.85 | 0.642 | 0.478 | 0.646 | 0.701 | 0.704 | 0.617 |
| RNA degradation | 0.856 | 0.787 | 0.743 | 0.73 | 0.646 | 0.703 | 0.656 | 0.653 | 0.645 | 0.675 |
| RNA transport | 0.859 | 0.82 | 0.794 | 0.855 | 0.735 | 0.733 | 0.727 | 0.636 | 0.731 | 0.683 |
| Salivary secretion | 0.739 | 0.722 | 0.762 | 0.721 | 0.64 | 0.654 | 0.72 | 0.644 | 0.67 | 0.607 |
| Shigellosis | 0.724 | 0.768 | 0.77 | 0.697 | 0.647 | 0.562 | 0.666 | 0.618 | 0.759 | 0.731 |
| Small cell lung cancer | 0.701 | 0.66 | 0.652 | 0.65 | 0.64 | 0.71 | 0.67 | 0.6 | 0.671 | 0.666 |
| SNARE interactions in vesicular transport | 0.85 | 0.877 | 0.856 | 0.827 | 0.664 | 0.751 | 0.789 | 0.677 | 0.774 | 0.644 |
| Spliceosome | 0.938 | 0.946 | 0.868 | 0.961 | 0.751 | 0.821 | 0.907 | 0.769 | 0.854 | 0.771 |
| Staphylococcus aureus infection | 0.899 | 0.976 | 0.983 | 0.979 | 0.853 | 0.831 | 0.902 | 0.959 | 0.971 | 0.955 |
| Starch and sucrose metabolism | 0.816 | 0.753 | 0.691 | 0.662 | 0.65 | 0.723 | 0.766 | 0.674 | 0.751 | 0.722 |
| Systemic lupus erythematosus | 0.899 | 0.933 | 0.82 | 0.913 | 0.765 | 0.786 | 0.934 | 0.979 | 0.867 | 0.834 |
| T cell receptor signaling pathway | 0.822 | 0.742 | 0.771 | 0.829 | 0.685 | 0.644 | 0.709 | 0.709 | 0.689 | 0.692 |
| TGF-beta signaling pathway | 0.87 | 0.736 | 0.741 | 0.881 | 0.607 | 0.63 | 0.772 | 0.565 | 0.837 | 0.678 |
| Tight junction | 0.778 | 0.686 | 0.682 | 0.618 | 0.594 | 0.551 | 0.624 | 0.603 | 0.613 | 0.642 |
| Toll-like receptor signaling pathway | 0.671 | 0.677 | 0.619 | 0.694 | 0.617 | 0.574 | 0.631 | 0.607 | 0.65 | 0.542 |
| Toxoplasmosis | 0.757 | 0.747 | 0.702 | 0.812 | 0.565 | 0.604 | 0.675 | 0.611 | 0.7 | 0.673 |
| Type I diabetes mellitus | 0.929 | 0.93 | 0.931 | 0.96 | 0.768 | 0.713 | 0.96 | 0.902 | 0.963 | 0.905 |
| Ubiquitin mediated proteolysis | 0.775 | 0.726 | 0.746 | 0.81 | 0.599 | 0.63 | 0.653 | 0.608 | 0.559 | 0.661 |
| Vascular smooth muscle contraction | 0.873 | 0.791 | 0.748 | 0.757 | 0.718 | 0.748 | 0.699 | 0.61 | 0.759 | 0.627 |
| Vasopressin-regulated water reabsorption | 0.899 | 0.829 | 0.841 | 0.797 | 0.619 | 0.657 | 0.706 | 0.586 | 0.817 | 0.666 |
| VEGF signaling pathway | 0.721 | 0.689 | 0.726 | 0.656 | 0.525 | 0.552 | 0.637 | 0.678 | 0.553 | 0.551 |
| Vibrio cholerae infection | 0.909 | 0.841 | 0.844 | 0.82 | 0.67 | 0.647 | 0.62 | 0.718 | 0.731 | 0.621 |
| Viral myocarditis | 0.695 | 0.763 | 0.767 | 0.806 | 0.659 | 0.598 | 0.673 | 0.776 | 0.763 | 0.834 |
| Wnt signaling pathway | 0.729 | 0.721 | 0.738 | 0.645 | 0.553 | 0.527 | 0.74 | 0.601 | 0.641 | 0.624 |

The rank boxplots below summarize the relative performance of the data tables in the functional prediction analysis. For each functional category, a rank is assigned to each data table based on its AUROC compared to the other data tables, where the best functional prediction rank is 1 and the poorest rank is the number of data tables.

Comparison of each protein (RNA) data table to a designated RNA (protein) data table is also summarized in the scatter plots below. For each point, the AUROC for a given category in the RNA data is plotted on the X-axis whereas the corresponding AUROC in the protein data table is plotted on the Y-axis. The number of categories for which the protein data table outperforms the RNA data table (AUROC(protein) > 1.1 \* AUROC (RNA); red dots) and vice versa (AUROC(RNA) > 1.1 \* AUROC (protein); blue dots) are also shown.

FP-RefRatioFP-RefRatio-MDFP-RefRatio-Weighted-MDFP-VirtualRatio-MDMQ-NoNormMQ-NoNorm-MDMQ-RefRatio-MDMQ-RefRatio-Weighted-MDMQ-VirtualRatio-MD

#### 6.3 PCA with sample class annotation

Another approach for assessing how well each data table can distinguish between classes is to determine how well each class can be separated by principal component analysis (PCA). In PCA score plots for each data table below, each point is a sample that is colored by class and that has a shape reflecting the batch. For a given sample, the PC2 score is plotted on the Y-axis whereas the PC1 score is plotted on the X-axis. Ellipses highlighting clusters of samples in each class are colored by corresponding class, and the separation between these ellipses indicates how well the variances captured by the first two PCs can distinguish between samples from different classes.

FP-RefRatio-MDFP-RefRatio-Weighted-MDFP-RefRatioFP-VirtualRatio-MDMQ-NoNorm-MDMQ-NoNormMQ-RefRatio-MDMQ-RefRatio-Weighted-MDMQ-VirtualRatio-MD

#### 6.4 Unsupervised clustering

Unsupervised hierarchical clustering can reveal patterns in the data (clusters of genes or samples that behave more similarly to each other than to other genes or samples). Each heatmap below shows the results of hierarchical clustering for a given data table using `ComplexHeatmap`. Genes/proteins are in rows, while samples are in columns and labeled with corresponding class to visualize any potential associations between classes and clusters.

FP-RefRatio-MDFP-RefRatio-Weighted-MDFP-RefRatioFP-VirtualRatio-MDMQ-NoNorm-MDMQ-NoNormMQ-RefRatio-MDMQ-RefRatio-Weighted-MDMQ-VirtualRatio-MD

### 7 Multi-omics concordance

The concordance between the protein data and RNA data can be used to assess data quality when both RNA and protein data tables are available. Here, we evaluate gene- and sample-wise correlations between the protein and RNA data tables.

#### 7.1 Gene-wise mRNA-protein correlation

The table below shows the number of genes with measurements (n) in each data table as well as the median of all gene-wise Spearman correlations between mRNA and protein measurements. The columns n5, n6, n7 and n8 show the number of genes with correlation greater than 0.5, 0.6, 0.7 and 0.8, respectively.

| data table | n | n5 | n6 | n7 | n8 | gene\_wise\_cor |
| --- | --- | --- | --- | --- | --- | --- |
| FP-RefRatio-MD | 9034 | 3372 | 2217 | 1204 | 467 | 0.4050 |
| FP-RefRatio-Weighted-MD | 9034 | 3230 | 2107 | 1137 | 432 | 0.3939 |
| FP-RefRatio | 9034 | 1586 | 799 | 319 | 79 | 0.2792 |
| FP-VirtualRatio-MD | 9034 | 2921 | 1735 | 806 | 202 | 0.3720 |
| MQ-NoNorm-MD | 8043 | 936 | 459 | 189 | 58 | 0.2127 |
| MQ-NoNorm | 8043 | 521 | 206 | 71 | 11 | 0.1875 |
| MQ-RefRatio-MD | 8042 | 2993 | 1972 | 1062 | 395 | 0.4071 |
| MQ-RefRatio-Weighted-MD | 8042 | 1244 | 617 | 196 | 27 | 0.2625 |
| MQ-VirtualRatio-MD | 8043 | 2443 | 1428 | 588 | 138 | 0.3667 |

Spearman correlation results are also shown for each gene/protein in the boxplot below.

Another way to visualize the differences between the distributions of all gene-wise RNA-protein correlations is with the cumulative distribution function (CDF) plot shown below. Here each line shows the cumulative distribution for the gene-wise correlations. The further the distribution function is shifted to the right, the more highly correlated the RNA-protein data is.

The histograms below provide another way to visualize the distribution of correlations for each protein (or RNA) data table with the RNA (or protein) data. Here the bars showing binned frequencies of positive correlations are in red, while negative correlations are shown in the blue bins, and summary statistics are also provided.

FP-RefRatio-MDFP-RefRatio-Weighted-MDFP-RefRatioFP-VirtualRatio-MDMQ-NoNorm-MDMQ-NoNormMQ-RefRatio-MDMQ-RefRatio-Weighted-MDMQ-VirtualRatio-MD

#### 7.2 Sample-wise mRNA-protein correlation

Sample-wise RNA-protein correlations are summarized in the table below as the median of Spearman correlations for matched protein and RNA data from all pairs of samples for each data table, while the violin plots below show the distributions of these correlations for each data table.

| data table | sample\_wise\_cor |
| --- | --- |
| FP-RefRatio | 0.4641 |
| FP-RefRatio-MD | 0.4641 |
| FP-RefRatio-Weighted-MD | 0.4664 |
| FP-VirtualRatio-MD | 0.4594 |
| MQ-NoNorm | 0.4148 |
| MQ-NoNorm-MD | 0.4148 |
| MQ-RefRatio-MD | 0.1856 |
| MQ-RefRatio-Weighted-MD | 0.3047 |
| MQ-VirtualRatio-MD | 0.1693 |
