## Supplementary Figures for "Analysis of isobaric quantitative proteomic data using TMT-Integrator and FragPipe computational platform"

**Supplement to: “Analysis of data from isobaric labeling-based proteomic experiments using TMT-Integrator and FragPipe computational platform”**

Hui-Yin Chang^1,2#^, Yamei Deng^1#,^ Ruohong Li^1^, Dmitry Avtonomov^1^, Bo Wen^3^, Sarah E. Haynes^1^, Felipe da Veiga Leprevost^1^, Bing Zhang^3^, Fengchao Yu^1^, Alexey I. Nesvizhskii^1,4,*^

^1^ Department of Pathology, University of Michigan, Ann Arbor, Michigan, United States

^2^ Department of Biomedical Sciences and Engineering, National Central University, Taiwan

^3^ Baylor College of Medicine, Houston, Texas, United States

^4^ Department of Computational Medicine and Bioinformatics, University of Michigan, Ann Arbor, Michigan, United States

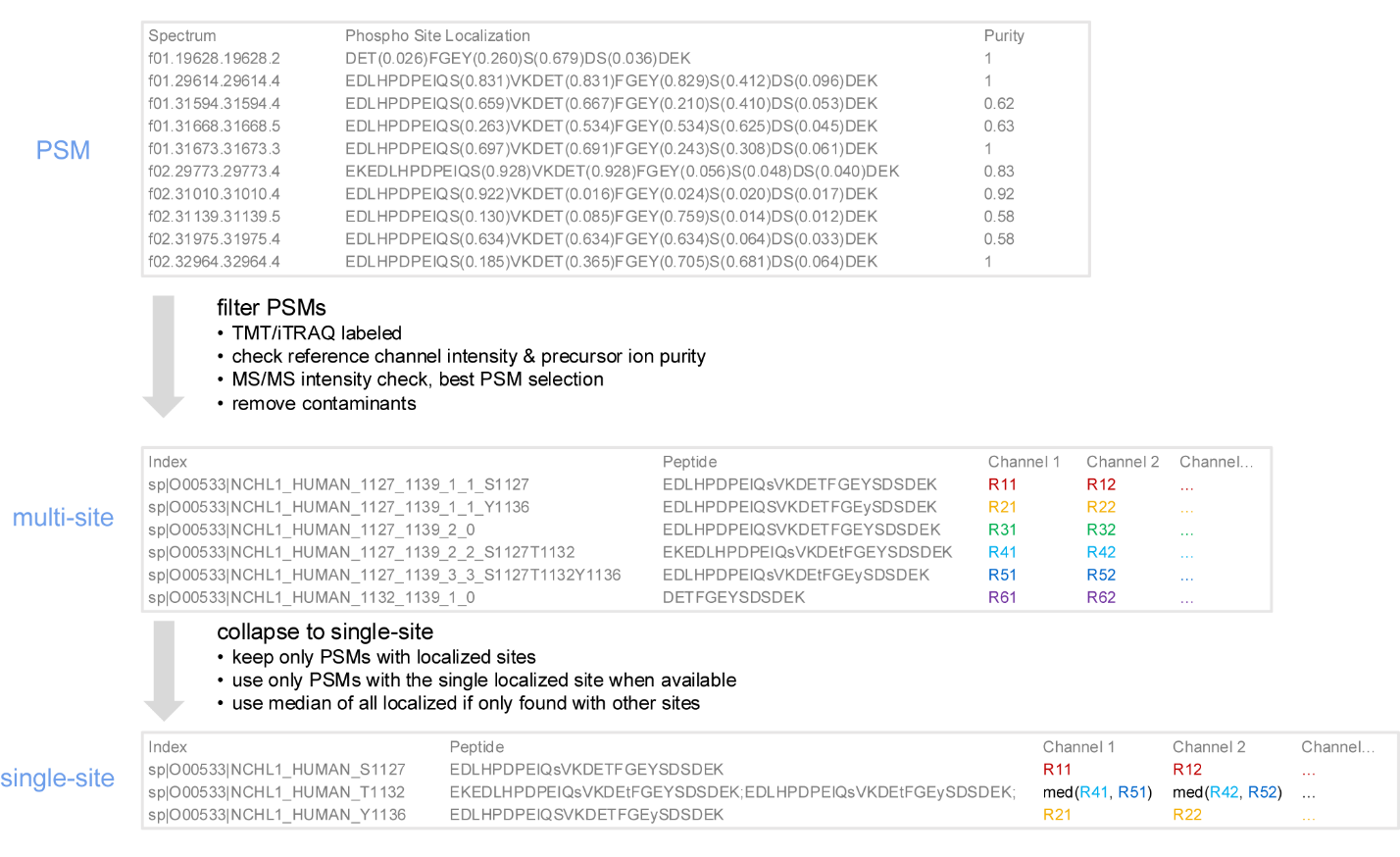

**Supplementary Figure 1. Example illustrating the detailed process of generating single-site ratio table, starting from the PSM list to multi-site ratios and finally to single-site ratios.** Ratio values are denoted as colored text in the channel columns.

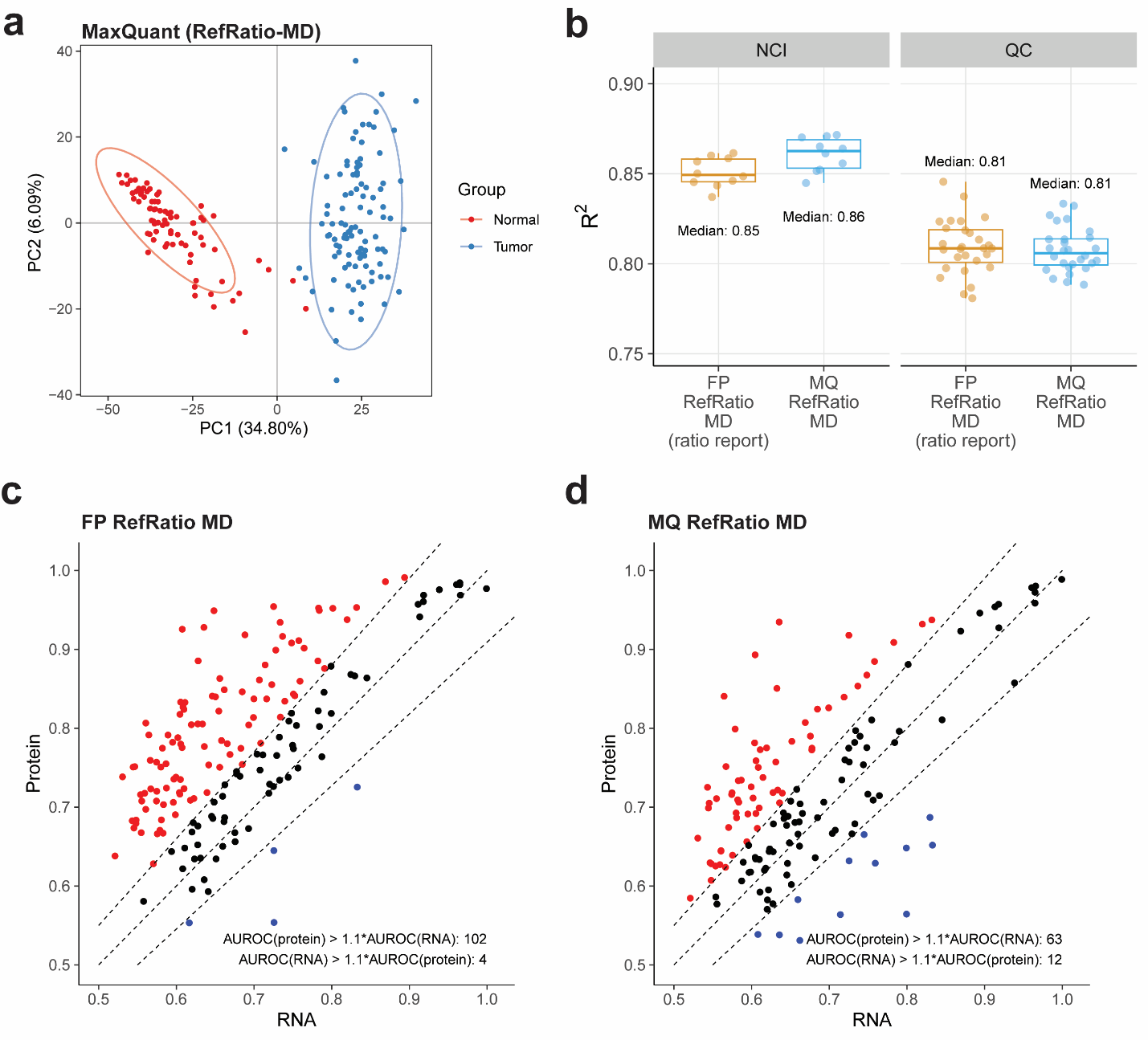

**Supplementary Figure 2. Performance evaluations on ccRCC whole proteome dataset.** **(a)** PCA plot of MaxQuant median-centered gene quantification from tumor and normal samples. **(b)** Boxplots showing the R-squared value distributions for protein abundance correlation among replicate runs of the NCI and QC samples using ratio-to-reference data, with median correlation coefficients (R-square) labeled for each method. FP represents FragPipe, MQ represents MaxQuant, RefRatio represents ratio-to-reference normalization with a real reference, VirtualRatio represents ratio-to-reference normalization with a virtual reference and MD represents the use of median-centering normalization. **(c)** Scatter plot showing the difference in KEGG pathway membership predictions (AUROC values) by OmicsEV using the protein data table from FragPipe (RefRatio MD) along with the RNA data table. Each dot represents a gene used in the predictions: red dots indicate genes where AUROC values for protein data exceed those for RNA data by a specified threshold, blue dots indicate genes where AUROC values for RNA data exceed those for protein data by the threshold, and black dots indicate genes with no noticeable difference in AUROC values between protein and RNA data. **(d)** Same as **(c)** for the protein data table from MaxQuant (RefRatio MD).

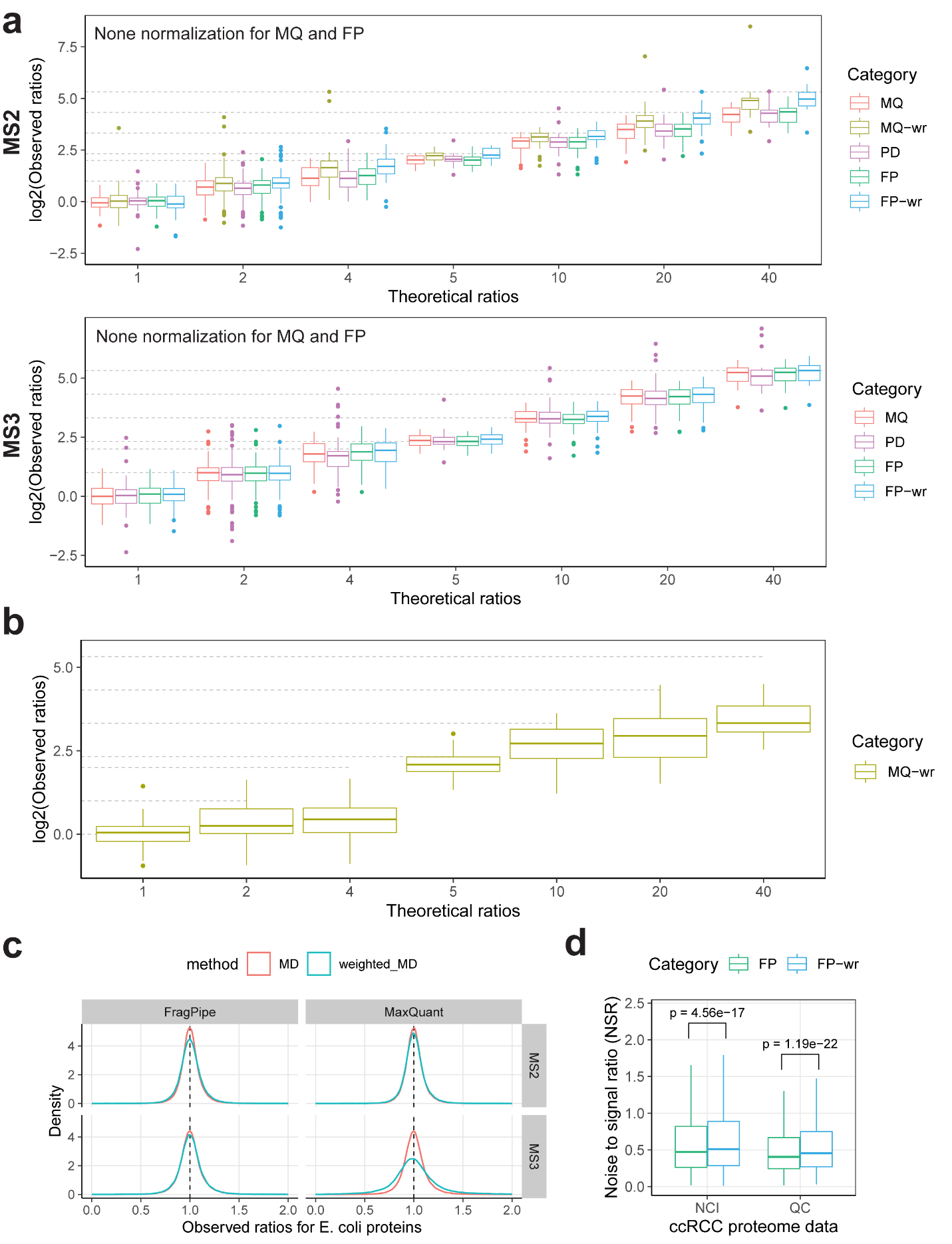

**Supplementary Figure 3. Performance evaluations on the spiked-in dataset. (a)** Boxplots showing the observed ratio distribution of 12 spiked-in proteins compared to theoretical ratios (grey dashed lines) using non-median-centered MS2 and MS3 data across different methods. **(b)** Boxplots showing the observed ratio distribution of 12 spiked-in proteins compared to theoretical ratios (grey dashed lines) in MS2 and MS3 data for MaxQuant weighted method. **(c)** Comparison of two ratio integration approaches (conventional median ratio method and weighted ratio method) based on the observed ratio distributions for E. coli proteins for FragPipe and MaxQuant. Density plots show the observed ratio distributions of E. coli proteins using MS2 and MS3 data. Line colors represent different ratio integration methods. **(d)** Comparison of two ratio integration approaches in FragPipe based on protein noise-to-signal ratio (NSR) levels in ccRCC whole proteome data. Boxplots depict NSR distributions in NCI and QC channels, with significance p-values from t-test labeled.

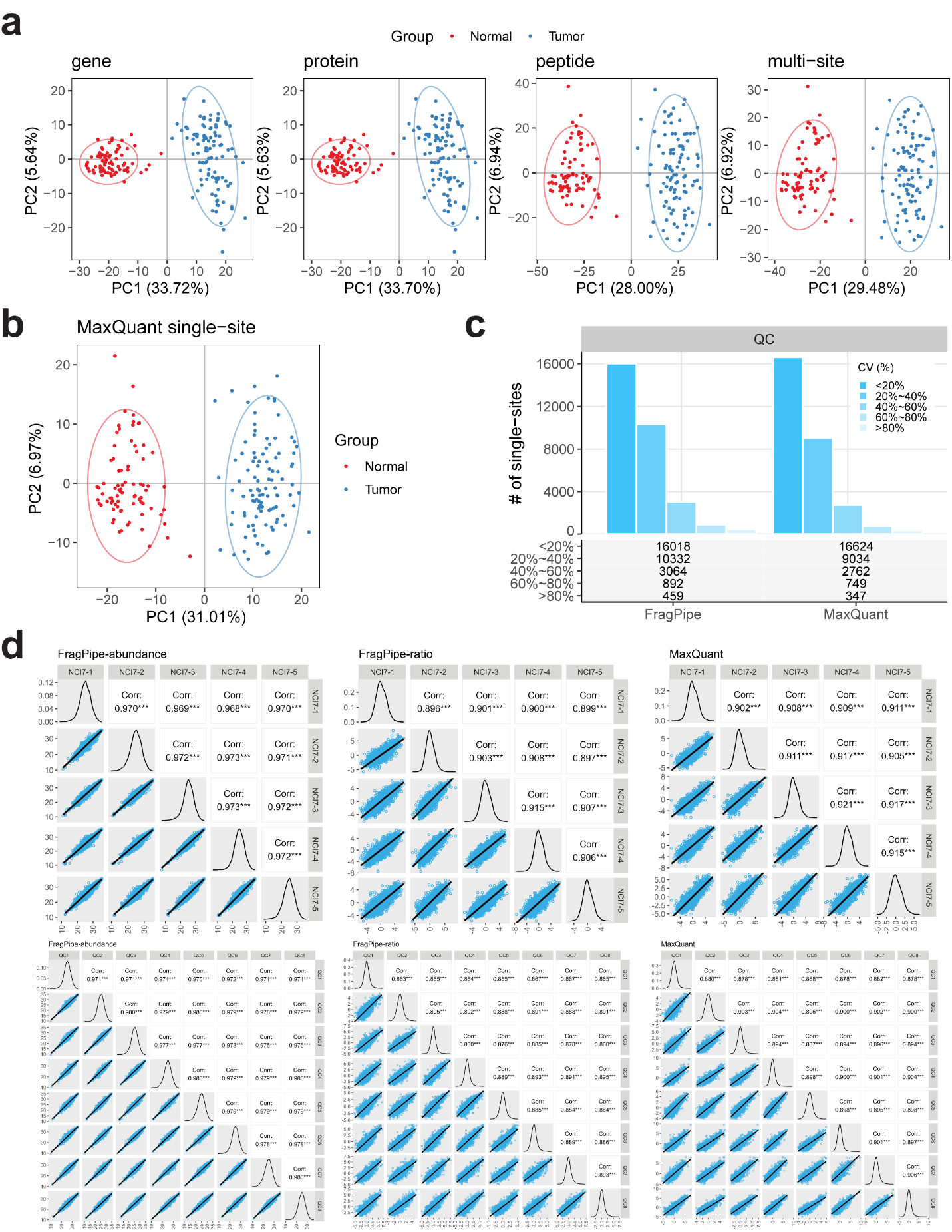

**Supplementary Figure 4. Performance evaluations on the ccRCC phosphorylation-enriched dataset. (a)** PCA plots of TMT-Integrator median-centered data for gene, protein, peptide and multi-site levels from tumor and normal samples. **(b)** PCA plot of MaxQuant median-centered single-site data from tumor and normal samples. **(c)** Comparison of single-site variation (CV) distributions in QC channels between FragPipe and MaxQuant, with bars representing the number of single-sites in each CV group. **(d)** Evaluation of quantification consistency across FragPipe abundance and ratio single-site reports and MaxQuant single-site ratio data in NCI and QC channels. Each subplot is labeled with the data type in the top left. The lower triangular shows linear model fitting for each sample pair, the diagonal displays density plots of overall abundance or ratio distributions for each sample, and the upper triangular presents Pearson correlation test results, with correlation coefficients and significance (*) indicated.
